# Single Photon smFRET. III. Application to Pulsed Illumination

**DOI:** 10.1101/2022.07.20.500892

**Authors:** Matthew Safar, Ayush Saurabh, Bidyut Sarkar, Mohamadreza Fazel, Kunihiko Ishii, Tahei Tahara, Ioannis Sgouralis, Steve Pressé

**Affiliations:** Center for Biological Physics, Arizona State University, Tempe, AZ, USA; Department of Mathematics and Statistical Science, Arizona State University, Tempe, AZ, USA; Department of Physics, Arizona State University, Tempe, AZ, USA; Molecular Spectroscopy Laboratory, RIKEN, 2-1 Hirosawa, Wako, Saitama 351-0198, Japan; Ultrafast Spectroscopy Research Team, RIKEN Center for Advanced Photonics (RAP), 2-1 Hirosawa, Wako, Saitama 351-0198, Japan; Department of Mathematics, University of Tennessee Knoxville, Knoxville, TN, USA; School of Molecular Sciences, Arizona State University, Phoenix, AZ, USA

## Abstract

Förster resonance energy transfer (FRET) using pulsed illumination has been pivotal in leveraging lifetime information in FRET analysis. However, there remain major challenges in quantitative single photon, single molecule FRET (smFRET) data analysis under pulsed illumination including: 1) simultaneously deducing kinetics and number of system states; 2) providing uncertainties over estimates, particularly uncertainty over the number of system states; 3) taking into account detector noise sources such as crosstalk, and the instrument response function contributing to uncertainty; in addition to 4) other experimental noise sources such as background. Here, we implement the Bayesian nonparametric framework described in the first companion manuscript that addresses all aforementioned issues in smFRET data analysis specialized for the case of pulsed illumination. Furthermore, we apply our method to both synthetic as well as experimental data acquired using Holliday junctions.

**Why It Matters:** In the first companion manuscript of this series, we developed new methods to analyze noisy smFRET data. These methods eliminate the requirement of *a priori* specifying the dimensionality of the physical model describing a molecular complex’s kinetics. Here, we apply these methods to experimentally obtained datasets with samples illuminated by laser pulses at regular time intervals. In particular, we study conformational dynamics of Holliday junctions.

## 1 Terminology Convention

To be consistent throughout our three part manuscript, we precisely define some terms as follows

1. a macromolecular complex under study is always referred to as a *system*,
2. the configurations through which a system transitions are termed *system states*, typically labeled using *σ*,
3. FRET dyes undergo quantum mechanical transitions between *photophysical states*, typically labeled using *ψ*,
4. a system-FRET combination is always referred to as a *composite*,
5. a composite undergoes transitions among its *superstates*, typically labeled using *ϕ*,
6. all transition rates are typically labeled using *λ*,
7. the symbol *N* is generally used to represent the total number of discretized time windows, typically labeled with *n*, and
8. the symbol *w_n_* is generally used to represent the observations in the *n*-th time window.

## 2 Introduction

Amongst the many fluorescence methods available [1–7], single molecule Förster resonance energy transfer (smFRET) has been useful in probing interactions and conformational changes on nanometer scales [8–12]. This is typically achieved by estimating FRET efficiencies (and system states) at all instants of an smFRET trace and subsequently estimating transition rates. Furthermore, among different FRET modalities, FRET efficiencies are most accurately determined under pulsed illumination [13–15], where the FRET dyes are illuminated by short laser bursts at known times.

Under this illumination procedure, photon arrival times are recorded with respect to the immediately preceding pulse, thereby facilitating accurate estimation of fluorescence lifetimes as well as FRET rates. As such, in this manuscript, we will focus on single photon smFRET analysis under pulsed illumination.

Under pulsed illumination, information on kinetic parameters present in smFRET data is traditionally learned by: binned photon methods thereby eliminating lifetime information altogether [16–18]; bulk correlative methods [19–21]; and single photon methods [14, 22, 23]. However, these methods are parametric, *i.e*., require fixing the number of system states *a priori*, and necessarily only learn system kinetics even though information on the number of system states is encoded in the data.

In this paper, we implement a general smFRET analysis framework presented in the Sec. 2.5.1 of the first companion manuscript [24] for the case of pulsed illumination to learn full distributions. In other words, probability distributions over parameters taking into account uncertainties from all existing sources such as crosstalk and background. These parameters include the system transition probabilities, and photophysical rates, that is, donor and acceptor relaxation and FRET rates, with special attention paid to uncertainty arising from sources such as inherent stochasticity in photon arrival times and detectors. As our main concern is deducing the number of system states using single photon arrivals while incorporating detector effects, we leverage the formalism of infinite hidden Markov models (iHMM) [25–30] within the Bayesian nonparametric (BNP) paradigm [25, 26, 31–38]. The iHMM framework assumes an *a priori* infinite number of system states with associated transition probabilities, where the number of system states warranted by input data is enumerated by those states most visited over the course of the system state trajectory.

Next, to benchmark our BNP-FRET sampler, we analyzed synthetic and experimental smFRET data acquired using a single confocal microscope with pulsed illumination optimized to excite donor dyes.

In particular, we employ a broad range of experimental data acquired from Holliday junctions (HJ) with an array of different kinetic rates due to varying buffer concentration of MgCl_2_ [39–42].

## 3 Forward Model and Inverse Strategy

In this section, we first briefly illustrate the adaptation of the general formalism described in our first companion manuscript [24] to the pulsed illumination case. Next, we present a specialized inference procedure. The details of the framework not provided herein can be found in the Supplementary Information.

As before, we consider a molecular complex labeled with a donor-acceptor FRET pair. As the molecular complex transitions through its *M_σ_* system states indexed by 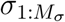, laser pulses (optimized to excite the donor) separated by time *τ* may excite either the donor or acceptor to drive transitions among the photophysical states, 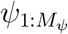, as defined in the first companion manuscript [24]. Such photophysical transitions lead to photon emissions that may be detected in either donor or acceptor channels. The set of *N* observations, *e.g*., photon arrival times, from *N* pulses are recorded as

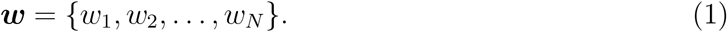

Here, each individual measurement is a pair 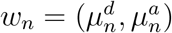, where 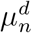 and 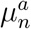 are the recorded arrival times (also known as microtimes) after the *n*-th pulse in both donor and acceptor channels, respectively. In cases where there is no photon detection, we denote the absent microtimes with 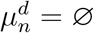 and 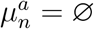 for donor and acceptor channels, respectively.

As is clear from Fig. 1, smFRET traces are inherently stochastic due to the nature of photon excitation, emission, and noise introduced by detector electronics. To analyze such stochastic systems, we begin with the most generic likelihood derived in Eq. 51 of the first companion manuscript [24]

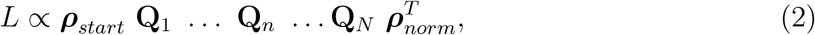

where ***ρ***_*start*_ is the initial probability vector for the system-FRET composite to be in one of *M* (= *M_ψ_* × *M_σ_*) superstates, and ***ρ***_*norm*_ is a vector that sums the elements of the propagated probability vector. Here, we recall that **Q**_*n*_ is the transition probability matrix between pulses *n* and *n* + 1, characterizing system-FRET composite transitions among superstates.

**Figure 1:**
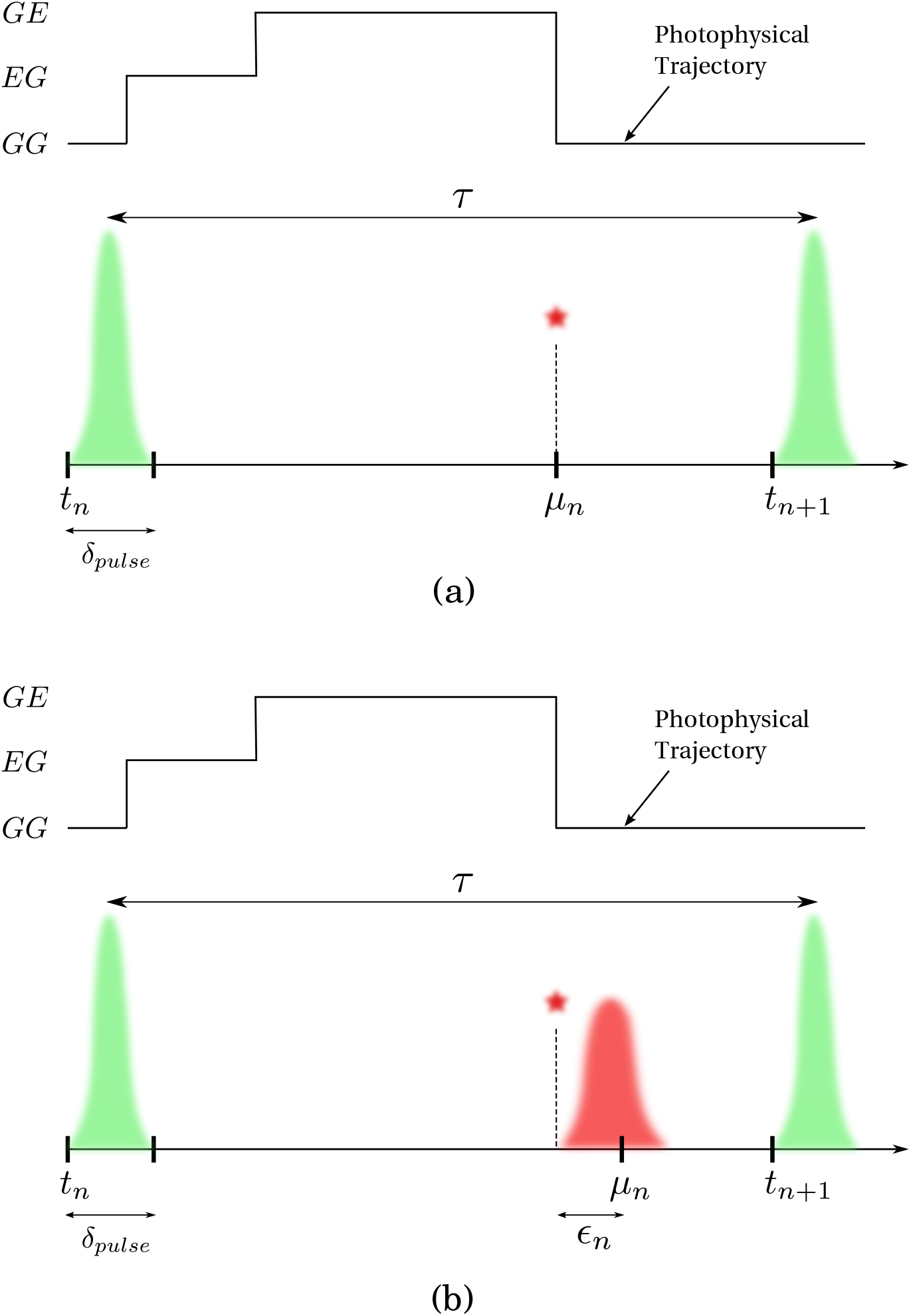
Events over a pulsed illumination experiment pulse window. Here, the beginning of the *n*-th interpulse window of size *τ* is marked by time *t_n_*. The FRET labels originally in state GG (donor and acceptor, respectively, in ground states) are excited by a high intensity burst (shown by the green) to the state EG (only donor excited) for a very short time *δ_pulse_*. If FRET occurs, the donor transfers its energy to the acceptor and resides in the ground state leaving the FRET labels in the GE state (only acceptor excited). The acceptor then emits a photon to be registered by the detector at microtime *μ_n_*. When using ideal detectors, the microtime is the same as the photon emission time as shown in panel (a). However, when the timing hardware has jitter (shown in red), a small delay *ϵ_n_* is added to the microtime as shown in panel (b). For convenience, we have reproduced this figure from our first companion manuscript [24].

The propagators **Q**_*n*_ above adopt different forms depending on whether a photon is detected or not during the associated period. Their most general forms are derived in Sec. 2.5.1 of the first companion manuscript [24]. However, these propagators involve computationally expensive integrals and thus we make a few approximations here as follows: 1) we assume that the system state remains the same over an interpulse period since typical system kinetic timescales (typically 1 ms or more) are much longer than interpulse periods (≈ 100 ns) [41, 43]; 2) the interpulse period (≈ 100 ns) is longer than the donor and acceptor lifetimes (≈ a few ns) [41, 43] such that they relax to the ground state before the next pulse. Furthermore, we will demonstrate a specialized sampling scheme under these physically motivated approximations.

The immediate implications of assumption (1) are that the system transitions may now, to a good approximation, only occur at the beginning of each pulse. Consequently, the evolution of the FRET pair between two consecutive pulses is now exclusively photophysical as the system state remains the same during interpulse times. As such, the system now evolves in equally spaced discrete time steps of size *τ* where the system state trajectory can be written as

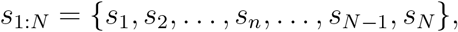

where *s_n_* is the system state between pulses *n* and *n* + 1. The stochastic evolution of the system states in such discrete steps is then determined by the transition probability matrix designated by **Π**_*σ*_. For example, in the simplest case of a molecular complex with two system states *σ*_1:2_, this matrix is computed as follows

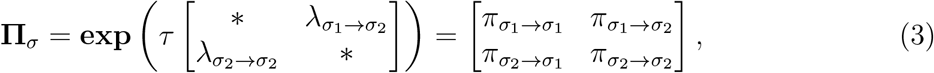

where the matrix in the exponential contains transition rates among the system states and the * represents the negative row sum.

Next, by assumption 2, we can further suppose that the fluorophores always start in the ground state at the beginning of every pulse. As a result, we treat pulses independently and write the probability of observation *w_n_* as

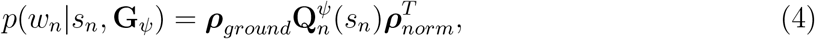

where ***ρ***_*ground*_ denotes the probability vector when the FRET pair is in the ground state at the beginning of each pulse, **G**_*ψ*_ is the generator matrix with only photophysical transition rates, and 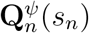 is the photophysical propagator for the *n*-th interpulse period.

We further organize the observation probabilities of Eq. 4 into a newly defined detection matrix 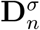 with its elements given by 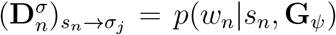. Here, we note that the index *j* does not appear on the right hand side because the system state does not change during an interpulse window resulting in the independence of observation probability from the next system state *s*_*n*+1_. The explicit formulas for the observation probabilities are provided in the Supplementary Information.

Now, using the matrix 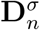, we define the reduced propagators for each interpulse period as

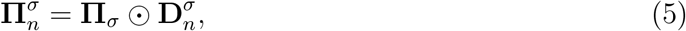

where ⊙ denotes element-by-element product.

Finally, using these simplified propagators, we can write the likelihood for a an smFRET trace under pulsed illumination as

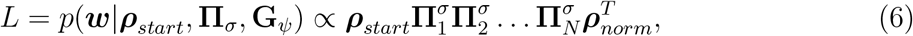

as also introduced in the Sec. 2.5.1 of the first companion manuscript [24]. This form of the likelihood is advantageous in that it allows empty pulses to be computed as a simple product, greatly reducing computational cost.

In the following, we first illustrate a parametric inference procedure assuming a given number of system states. We next generalize the procedure developed to the nonparametric case to deduce the number of system states along with the rest of parameters.

### 3.1 Inference Procedure: Parametric Sampler

Now, with the likelihood at hand, we construct the posterior as follows

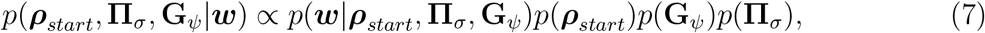

where we assume in the prior that the unknown parameters, including the initial probability vector, ***ρ***_*start*_, the photophysical transition rates in the generator matrix **G**_*ψ*_, and the transition probabilities among system states in propagator **Π**_*σ*_, are independent. Here, we can sample the set of unknowns using the above posterior with the Gibbs sampling procedure as described in the first companion manuscript Sec. 3.2.2. However, a computationally more convenient inference procedure that allows direct sampling is accomplished by writing the posterior of Eq. 8 as a marginalization (sum) over state trajectories as follows

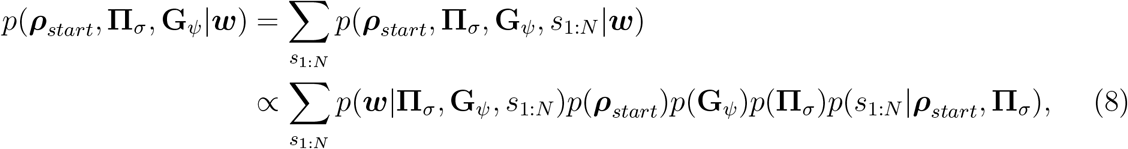

where *s*_1:*N*_ = {*s*_1_, *s*_2_ …, *s_N_*} denotes a system state trajectory. Now, we can use the non-marginal posterior

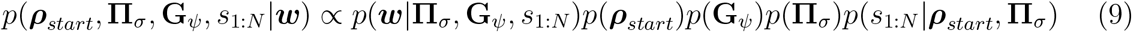

to sample the trajectory *s*_1:*N*_ which, in turn, allows direct sampling of the elements of propagator **Π**_*σ*_ described shortly. For priors on ***ρ***_*start*_ and rates in **G**_*ψ*_, we, respectively, use a Dirichlet and Gamma distributions similar to Eq. 65-66 of the first companion manuscript [24]. We sample the system state trajectory *s*_1:*N*_ by recursively sampling the states using a forward filtering backward sampling algorithm described in the Supplementary Information Sec. S4.3.

Finally, for each row in the propagator **Π**_*σ*_, we use a Dirichlet prior

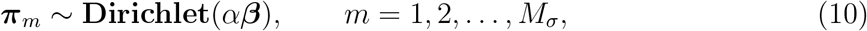

where *M_σ_* is the number of system states and ***π***_*m*_ denotes the *m*-th row of the propagator. Here, the hyperparameters *α* and ***β*** are, respectively, the concentration parameter and a vector of length *M_σ_* described in the first companion manuscript [24] Sec. 3.2.2. We can now directly generate samples for the transition probability vectors ***π***_*m*_ of length *M_σ_* via prior-likelihood conjugacy as (see Supplementary Information Sec. S4.3)

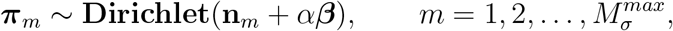

where the vector **n**_*m*_ collects the number of times each transition out of system state *σ_m_* occurs obtained using the system state trajectory.

After constructing the posterior, we can make inferences on the parameters by drawing samples from the posterior. However, as the resulting posterior has a nonanalytical form, it cannot be directly sampled. Therefore, we develop a Markov chain Monte Carlo sampling (MCMC) procedure [38, 44–48] to draw samples from the posterior.

Our MCMC scheme follows a Gibbs sampling technique sweeping through updates of the set of parameters in the following order: 1) photophysical transition rates including donor relaxation rates *λ_d_* (inverse of donor lifetime), acceptor relaxation rate *λ_a_* (inverse of acceptor lifetime), FRET rates 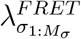 for each system state, and excitation rate (inverse of excitation probability *π_ex_*) using MH; 2) transition probabilities between system states, 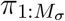 by directly drawing samples from the posterior; 3) the system states trajectory, 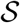, using forward backward sampling procedure [49]; and 4) the initial probabilities ***ρ***_*start*_ by taking direct samples. In the end, the chains of samples drawn can be used for subsequent numerical analysis.

### 3.2 Inference Procedure: Nonparametrics Sampler

The smFRET data analysis method illustrated above assumes a given number of system states, *M_σ_*. However, in many applications the number of system states is not specified *a priori*. Here, we describe a generalization of our parametric method to address this short-coming and estimate the number of system states simultaneously along with other unknown parameters.

We accomplish this by modifying our previously introduced parametric posterior as follows. First, we suppose an infinite number of system states (*M_σ_* → ∞) for the likelihood introduced previously and learn the transition matrix **Π**_*σ*_. The number of system states can then be interpreted as those appreciably visited over the course of the trajectory.

To incorporate this infinite system state space into our inference strategy, we leverage the iHMM [25, 26, 28–30] from the BNP repertoire, placing a hierarchical Dirichlet process prior over the infinite set of system states as described in the first companion manuscript, Sec. 3.2.2 [24]. However, as detailed in the first companion manuscript [24] Sec. 3.2.2 dealing with an infinite number of random variables, though feasible, is not computationally efficient and we approximate this infinite value with a large number 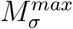, reducing our hierarchical Dirichlet process prior to

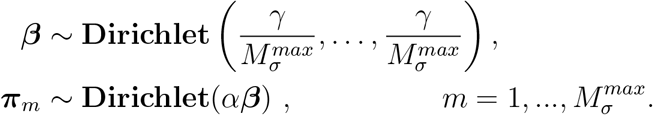

Here, ***β*** denotes the base probability vector of length 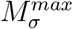 serving itself as a prior on the probability transition matrix **Π**_*σ*_, and ***π***_*m*_ is the *m*-th row of **Π**_*σ*_. Moreover, *γ* is a positive scalar hyperparameter of the Dirichlet process prior often chosen to be one. As such, we ascribe identical weights across the state space *a priori* for computational convenience [28, 29, 50].

Now, equipped with the nonparametric posterior, we proceed to simultaneously make inferences on transition probabilities, excited state escape rates, and the remaining parameters. To do so, we employ the Gibbs sampling scheme detailed in Sec. 3.2.2 on the first companion manuscript [24], except that we must now also sample the system state trajectory *s*_1:*N*_. More details on the overall sampling scheme are found in the Supplementary Information in section S4.

## 4 Results

The main objective of our method is to learn full distributions over: 1) transition probabilities among 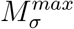 system states determining, in turn, the corresponding system transition rates and the effective number of system states; 2) photophysical transition rates including FRET rates 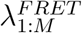, and fluorophores’ relaxation rates (inverse of lifetimes) *λ_a_* and *λ_d_*.

To sample from distributions over these parameters, the BNP-FRET sampler requires input data comprised of photon arrival time traces from both donor and acceptor channels as well as a set of precalibrated input parameters including: camera effects such as crosstalk matrix and detection efficiency (see Sec. 2.4 and Example V of the first companion manuscript [24]); background emission (see Sec. 2.6 of the first companion manuscript and Supplementary Information Sec. S2.4); and the IRF (see Sec. 2.5 of the first companion manuscript [24] and Supplementary Information Sec. S2.3).

Here, we first show that our method samples posteriors over a set of parameters employing realistic synthetic data generated using the Gillespie algorithm [51] to simulate system and photophysical transitions while incorporating detector artefacts such as crosstalk (see Sec. 2.8 of the first companion manuscript [24]). The list of parameters used in data generation for all the figures is provided in Supplementary Information Sec. S6. Furthermore, prior hyperparameters used in the analysis of synthetic and experimental data are listed in the Supplementary Information Sec. S3.

We first show that our method works for the simplest case of slow transitions compared to the interpulse period (25 ns) with two system states using synthetic data, see Fig. 2. Next, we proceed to tackle more challenging synthetic data with three system states and higher transition rates Fig. 3. We show that our nonparametric algorithm correctly infers system transition probabilities and thus the number of system states; see Fig. 3.

**Figure 2:**
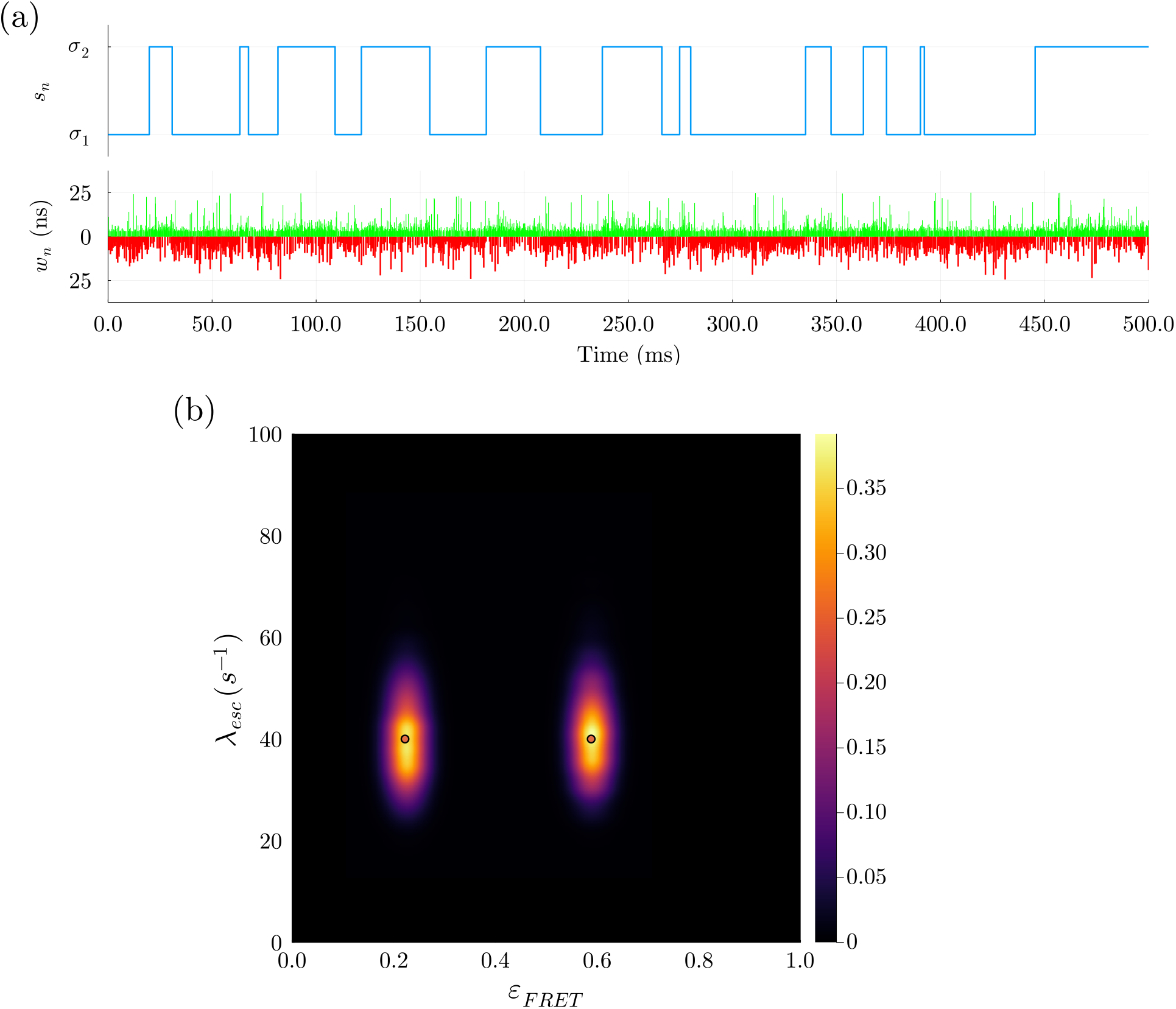
Analysis on synthetic data for a system with two system states. In panel (a), we show a section of synthetic data produced with the values in Supplementary Information Table S2. Furthermore, the system state trajectory is shown in blue. Below this, the arrival times of donor and acceptor photons 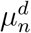 and 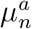 are shown in green and red, respectively. In panel (b), the ground truth is shown with red dots corresponding to escape rate of 40 s^−1^ and FRET efficiencies of 0.22 and 0.59. From our MAP estimate, we plot the bivariate distribution over escape rates *λ_esc_* and FRET efficiencies *ε_FRET_*. As seen, the BNP-FRET sampler clearly distinguishes two system states and locates ground truth values for the associated escape rates with values 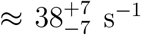 and 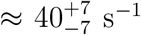, and FRET efficiencies 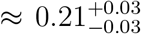 and 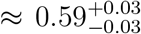. We have smoothed the distributions using kernel density estimation (KDE) for illustration purposes only.

**Figure 3:**
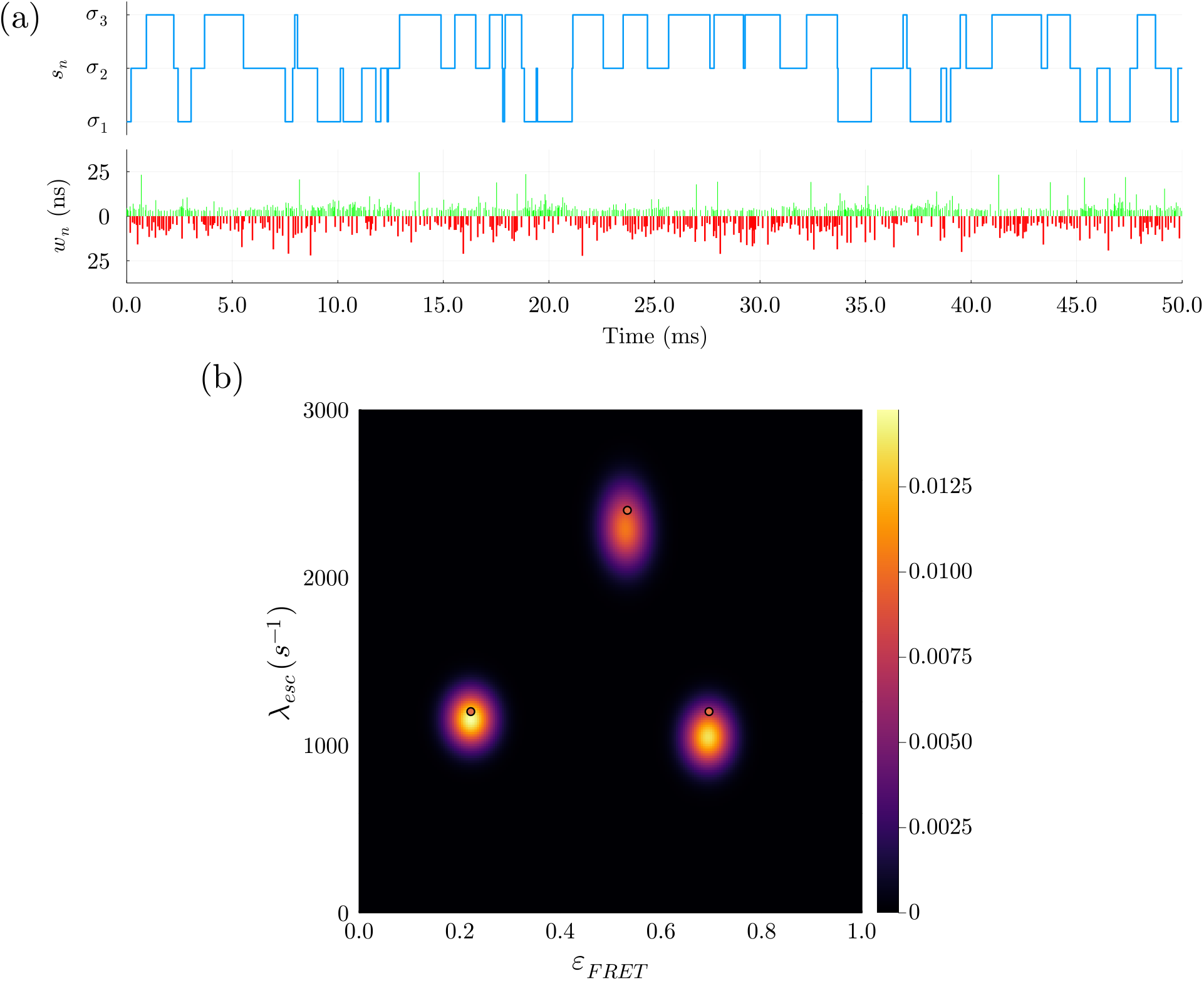
Analysis on synthetic data for three system states. In panel (a) we have a section of synthetic data produced with the values from Supplementary Information Table S3. The system state trajectory is seen in blue. Below this, the arrival times of donor and acceptor photons 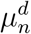 and 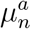 are shown in green and red, respectively. In panel (b), the red dots show ground truths corresponding to escape rates of 1200 s^−1^, 2400 s^−1^, and 1200 s^−1^ and FRET efficiencies of 0.22, 0.53 and 0.7. From our MAP estimate, we plot the distribution over escape rates *λ_esc_* and FRET efficiencies *ε_FRET_*. The MAP estimate clearly shows three system states with escape rates of 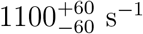, 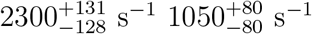.

After demonstrating the performance of our method using synthetic data, we use experimental data to investigate the kinetics of HJs under different MgCl_2_ concentrations in buffer; see Fig. 4.

**Figure 4:**
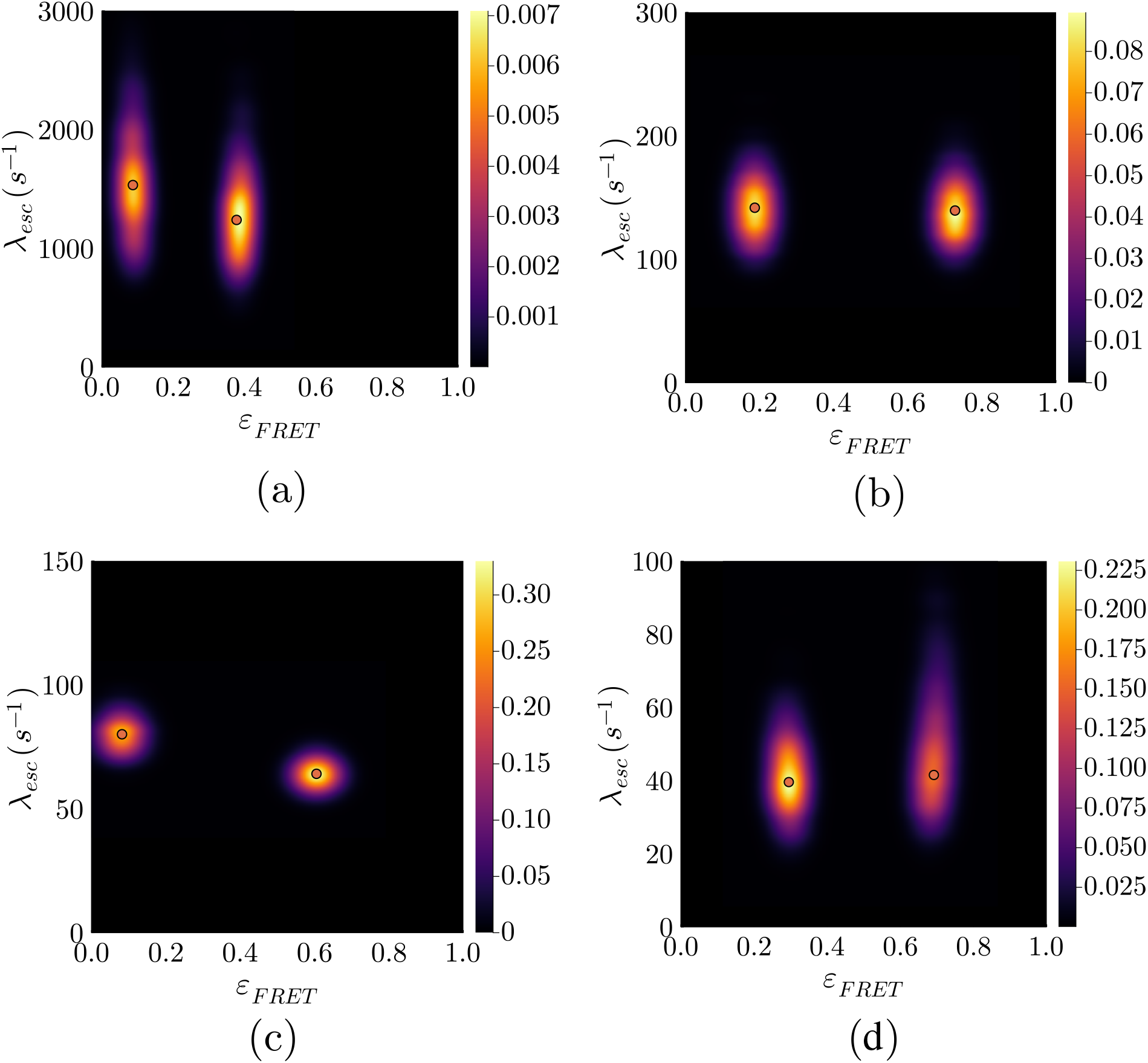
The bivariate posterior for the conformational transition rates *λ_esc_* and FRET efficiencies *ε_FRET_* for experimental data acquired in the presence of different HJ concentrations. Here, we show our bivariate posteriors where red dots show MAP estimates. In panel (a), we show the posterior for a sample with 1 mm MgCl_2_. We report escape rates of 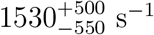 and 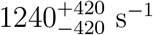 in this case. The posterior for a sample with 3 mm MgCl_2_ is shown in panel (b). We report escape rates of 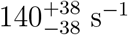 and 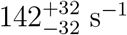 for this case. In panel (c), we show our posterior for a sample with 5 mm MgCl_2_. Here, we report escape rates of 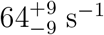 and 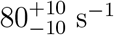. The posterior in panel (d) is for a sample with 10 mm MgCl_2_. We report escape rates of 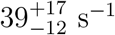 and 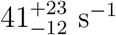.

### 4.1 Simulated Data Analysis

To help validate BNPs on smFRET single photon data, we start with a simple case of a two state system and select kinetics similar to those of the experimental data sets, *c.f*., the HJ in 10 mm MgCl_2_, with escape rates of 40 s^−1^ for both system states [52]. The generated system state trajectory and photon traces over a period of 500 ms from both channels are shown in Fig. 2 (a).

Fig. 2 (b) shows the bivariate posterior distribution over FRET efficiencies, *ε_FRET_*, defined as *ε_FRET_* = *λ_FRET_*/(*λ_FRET_* + *λ_d_*), and system escape rates, *i.e*., obtained by computing logarithm of the propagator matrix, with two peaks corresponding to two system states most visited by the sampler. Furthermore, the ground truths, designated by red dots, fall within the posterior with a relative error of less than 3% from the posterior modes. The results for the remaining parameters, including donor and acceptor transition rates, FRET transition rates and system transition probabilities, are presented in Supplementary Information Sec. S7.

To showcase the critical role played by BNPs, we also consider the more difficult case of a sample with three system states and faster system state kinetics ranging over 1200-2500 *s*^−1^. We do so by simulating photon traces in both donor and acceptor channels over a period of ~150 ms. A 50 ms section of the synthetic photon trace is shown in Fig 3(a).

Using direct photon arrivals from the generated photon trace, we find that the most probable system state trajectories sampled by BNP-FRET visit the correct number of system states, as shown in Fig 3(b), while inferring all other parameters. Furthermore, the BNP-FRET sampler estimates the system transition rates and thus the escape rates (*i.e*., sum of transition rates out of a given state) where the ground truth escape rates differ from the posterior peaks by a relative average error of less than 8%. The results for the remaining parameters are provided in the Supplementary Information Sec. S7.

### 4.2 Experimental Data Analysis: Holliday Junction (HJ)

In this section, we benchmark our method over a wide range of kinetic rates employing experimental data acquired using HJ under varying buffer MgCl_2_ concentrations [15, 52].

HJs are four-way double-helical DNA junctions existing in various structural configurations [41, 53, 54]. When not interacting with multivalent metal ions, electrostatic repulsion between negatively charged phosphate groups of the four helical arms forces HJs to assume a wide configuration where the arms lie along the two diagonals of a square. However, in the presence of ions, such as Mg^2+^, interaction with the phosphate groups results in electrostatic screening. This reduced repulsion induces transitions to what is believed to be primarily two compact stacked configurations/conformations. The transitions between both conformations necessitates passing through the intermediate open configuration. Since, at high ion concentrations, displacing ions away from the phosphate group becomes increasingly difficult, in this scenario we anticipate smaller transition rates between both conformations.

The HJ kinetic rates have been studied using both fluorescence lifetime correlation spectroscopy (FLCS) [15] and HMM analysis [55] on diffusing HJs assuming *a priori* a pair of high and low FRET system states. As expected, these previous studies show kinetic rates decreasing with increasing MgCl_2_ concentrations [41, 43] and correspondingly longer dwells.

Here our method, free from averaging and binning otherwise common in HMM analysis, is particularly well-suited to learn the rapid kinetics at low Mg^2+^ concentrations. We apply our BNP-FRET to data acquired from HJs at 1, 3, 5, and 10 mm MgCl_2_ concentrations, and sample the photophysical transition rates and the system transition probabilities.

The acquired bivariate posterior distributions over the FRET efficiencies and escape rates (computed via the logarithm of the system transition probability matrix **Π**_*σ*_) are presented in Fig. 4. Moreover, estimates for the other parameters can be found in the Supplementary Information Section S7. We note that our results are obtained on a single molecule basis with a photon budget of 10^4^ – 10^5^ photons.

For all four concentrations (see Fig. 4), our BNP-FRET sampler most frequently visited only two system states, while this was given as an input to the other analysis methods [15, 55]. Moreover, both escape rates are found to have similar values with an average of approximately 1400 s^−1^ (1 mM MgCl_2_), 140 s^−1^ (3 mM MgCl_2_), 72 s^−1^ (5 mM MgCl_2_), and 41 s^−1^ (10 mM MgCl_2_). These escape rates are in close agreement with values reported by FLCS and H2MM methods [15, 55] ≈ 1300 s^−1^ (1 mM MgCl_2_), ≈ 170 s^−1^ (3 mM MgCl_2_), ≈ 100 s^−1^ (5 mM MgCl_2_), ≈ 60 s^−1^ (10 mM MgCl_2_), which lie well within the bounds of our posteriors shown in Fig. 4 while simultaneously, and self-consistently, learning number of system states.

### 4.3 Experimental Data Acquisition

In this section, we describe the protocol for preparing the surface immobilized HJ sample labeled with a FRET pair and the procedure for recording smFRET traces from individual immobilized molecules. The sample preparation and recording of data follow previous work [56].

#### Sample preparation

The HJ used in this work consists of four DNA strands whose sequences are as follows

**R-strand**: 5’-CGA TGA GCA CCG CTC GGC TCA ACT GGC AGT CG-3’
**H-strand**: 5’-CAT CT<u>T</u> AGT AGC AGC GCG AGC GGT GCT CAT CG-3’
**X-strand**: 5’-biotin-TCTTT CGA CTG CCA GTT GAG CGC TTG CTA GGA GGA GC-3’
**B-strand**: 5’-GCT CC<u>T</u> CCT AGC AAG CCG CTG CTA CTA AGA TG-3’.

For surface immobilization, the X-strand was labeled with biotin at the 5’-end. For FRET measurements, the donor (ATTO-532) and acceptor (ATTO-647N) dyes were introduced into the H- and B-strands, respectively. In both cases, the dyes were labeled to thymine nucleotide at the 6th position from the 5’-ends of respective strands (shown as <u>T</u>). All DNA samples (labeled or unlabeled) were purchased from JBioS (Japan) in the HPLC purified form and were used without any further purification.

The HJ complex was prepared by mixing 1 mm solutions of R-, H-, B-, and X-strands in TN buffer (10 mm Tris-HCl with 50 mm NaCl, pH 8.0) at 3:2:3:3 molar ratio, annealing the mixture at 94 °C for 4 minutes, and gradually cooling it down (2-3 °C min^−1^) to room temperature (25 °C). For smFRET measurements, we used a sample chamber (Grace Bio-Labs SecureSeal, GBL621502) with biotin-PEG-SVA (biotin-poly(ethylene glycol)-succinimidyl valerate) coated coverslip. The chamber was first incubated with streptavidin (0.1 mg mL^−1^ in TN buffer) for 20 min. This was followed by washing the chamber with TN buffer (3 times) and injection of 1 nm HJ solution (with respect to its H-strand) for 3-10 seconds. After this incubation period, the chamber was rinsed with TN buffer (3 times) to remove unbound DNA and it was filled with TN buffer containing 1 mm (or 5 mm) MgCl_2_ and 2 mm Trolox for smFRET measurements.

#### smFRET measurements

The smFRET traces from individual HJs were recorded using a custom built confocal microscope (Nikon Eclipse Ti) equipped with the Perfect Focus System (PFS), a sample scanning piezo stage (Nano control B16-055), and a time correlated single photon counting (TCSPC) module (Becker and Hickl SPC-130EM).

The broadband light generated by a supercontinuum laser operating at 40 MHz (Fianium SC-400-4) was filtered with a bandpass filter (Semrock FF01-525/30) for exciting the donor dye, ATTO-532. This excitation light was introduced to the microscope using a single-mode optical fiber (Thorlabs P5-460B-PCAPC-1), and directed onto the sample using a dichroic mirror (Chroma ZT532/640rpc) and a water immersion objective lens (Nikon Plan Apo IR 60x, numerical aperture = 1.27).

The excitation light was focused onto the top surface of the coverslip and, during measurements, the focusing condition was maintained using the PFS. The fluorescence signals were collected by the same objective, passed through the dichroic mirror, and guided to the detection assembly (Thorlabs DFM1/M) using a multimode fiber (Thorlabs M50L02S-A). Note that this multimode fiber (core diameter: 50 μm) also acts as the confocal pinhole. In the detection assembly, the fluorescence signals from the donor and acceptor dyes were separated using a dichroic mirror (Chroma Technology ZT633rdc), filtered using bandpass filters (Chroma ET585/65m for donor, and Semrock FF02-685/40 for acceptor), and detected using separate hybrid detectors (Becker & Hickl HPM-100-40-C).

For each detected photon, its macrotime (absolute arrival time from the start of the measurement) was recorded with 25.2 ns resolution and its microtime (relative delay from the excitation pulse) was recorded with 6.1 ps resolution using the TCSPC module operating in time-tagging mode. A router (Becker and Hickl HRT-41) was used to process the signals from the donor and acceptor detectors.

For recording smFRET traces from individual HJs, we first imaged a 10 μm×3 μm area of the sample using the piezo stage by scanning it linearly at a speed of 1 μm s^−1^ in the X-direction and with an increment of 0.1 μm in the Y-direction. Individual HJs appeared as isolated bright spots in the image.

Next, we fitted the obtained donor and acceptor intensity images with multiple 2D Gaussian functions to determine the precise locations of individual HJs. Note that, during this image acquisition, the laser excitation power was kept to a minimum (~1 μW at the back aperture of the objective lens) to avoid photobleaching the dyes. In addition, we also employed an electronic shutter (Suruga Seiki, Japan) in the laser excitation path to control the sample excitation as required.

Using the precise locations of individual HJs obtained, we recorded 30 s long smFRET traces for each molecule by moving them to the center of the excitation beam using the piezo stage. For each trace, the laser excitation was blocked (using the shutter) for the first 5 seconds and was allowed to excite the sample for the remaining 25 seconds. Note that the smFRET traces were recorded using 40 μW laser excitation (at the back aperture of the objective lens) to maximize the fluorescence photons emitted from the dyes. We automated the process of acquiring smFRET traces from different molecules sequentially and executed it using a program written in-house on Igor Pro (Wavemetrics).

## 5 Discussion

The sensitivity of smFRET under pulsed illumination has been exploited to investigate many different molecular interactions and geometries [8–11, 57]. However, quantitative interpretation of smFRET data faces serious challenges including unknown number of system states and robust propagation of uncertainty from noise sources such as detectors and background. These challenges ultimately mitigate our ability to determine full distributions over all relevant unknowns and, traditionally, have resulted in data pre- or post-processing compromising the information otherwise encoded in the rawest form of data: single photon arrivals.

Here, we provide a general BNP framework for smFRET data analysis starting from single photon arrivals under a pulsed illumination setting. We simultaneously learn transition probabilities among system states as well as determine photophysical rates by incorporating existing sources of uncertainty such as background and crosstalk.

We benchmark our method using both experimental and simulated data. That is, we first show that our method correctly learns parameters for the simplest case with two system states and slow system transition rates. Moreover, we test our method on more challenging cases with more than two states using synthetic data and obtain correct estimations for the system state transition probabilities and thus the number of system states along with the remaining parameters of interest. To further assess our method’s performance, we analyzed experimental data from HJs suspended in solutions with a range of MgCl_2_ concentrations. These data were previously processed using other techniques assuming a fixed number of system states by binning photon arrival times [15].

Despite multiple advantages mentioned above for BNP-FRET, BNPs always come with an added computational cost as they take full advantage of information from single photon arrival times and all existing sources of uncertainty. For this version of our general BNP method simplified for pulsed illumination, we further reduced the computational complexity by grouping empty pulses together. Therefore, the computational complexity increased only linearly with the number of input photons as the photons are treated independently.

The method described in this paper assumes a Gaussian IRF. However, the developed framework is not limited to a specific form for the IRF and can be used for data collected using any type of IRF by modifying Eq. 4. Furthermore, the framework is flexible in accommodating different illumination techniques such as alternating color pulses, typically used to directly excite the acceptor fluorophores. This can be achieved by simple modification of the propagator 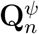 in Eq. 4. A future extension of this method could relax the assumption of a static sample by adding spatial dependence to the excitation rate as we explored in previous works [35, 48, 58]. This would allow our method to learn the dynamics of diffusing molecules, as well as their photophysical and system state transition rates.

## 6 Code Availability

The BNP-FRET software package is available on Github at https://github.com/LabPresse/BNP-FRET

## 7 Acknowledgments

We thank Dr. Zeliha Kilic, Weiqing Xu, and J Shephard Bryan IV for their contributions and insight into the project. We also thank Dr. Douglas Shepherd for providing insight into the workings of detectors and other experimental equipment and Irina Gopich for discussions on FRET likelihoods. T.T. acknowledges support from the JSPS KAKENHI (Grant No. 18H05265). S. P. acknowledges support from the NIH NIGMS (R01GM130745) for supporting early efforts in nonparametrics and NIH NIGMS (R01GM134426) for supporting single photon efforts.

