## Supplementary Information for "Single Photon smFRET. III. Application to Pulsed Illumination"

Last updated: November 1, 2022

#### Contents

**S1 Variables and Notation** **S1**

**S2 Likelihood for Pulsed Illumination** **S3**
S2.1 Excitation . . . . . . . . . . . . . . . . . . . . . . . . . . . . . . S4
S2.2 Photophysics . . . . . . . . . . . . . . . . . . . . . . . . . . . . . S5
S2.3 Instrument Response Function . . . . . . . . . . . . . . . . . . . . S7
S2.4 Background . . . . . . . . . . . . . . . . . . . . . . . . . . . . . . S8
S2.4.1 Laser background . . . . . . . . . . . . . . . . . . . . . . . . . S10
S2.4.2 Uniform background . . . . . . . . . . . . . . . . . . . . . . . . S11

**S3 Model Structure and Priors** **S13**
S3.1 Parametric Model . . . . . . . . . . . . . . . . . . . . . . . . . . . . . . S13
S3.2 Nonparametric Model . . . . . . . . . . . . . . . . . . . . . . . . . . . . S14

**S4 Sampling from the Posterior: Gibbs Algorithm Steps** **S15**
S4.1 Photophysical rates . . . . . . . . . . . . . . . . . . . . . . . . . . . . . S15
S4.2 Excitation Probability . . . . . . . . . . . . . . . . . . . . . . . . . . . S15
S4.3 System State Trajectory . . . . . . . . . . . . . . . . . . . . . . . . . . S16
S4.4 Transition probabilities . . . . . . . . . . . . . . . . . . . . . . . . . . S17
S4.5 Base Distribution . . . . . . . . . . . . . . . . . . . . . . . . . . . . . S17
S4.6 Optional Hyperparameters . . . . . . . . . . . . . . . . . . . . . . . . . S18
S4.6.1 Transition probabilities concentration hyperparameter . . . . . . . . . S18
S4.6.2 Sticky hyperparameter . . . . . . . . . . . . . . . . . . . . . . . . . S18
S4.6.3 Base distribution concentration hyperparameter . . . . . . . . . . . . S19

**S5 Estimation of pre-set Parameters** **S20**
S5.1 IRF . . . . . . . . . . . . . . . . . . . . . . . . . . . . . . . . . . . . S20
S5.2 Background . . . . . . . . . . . . . . . . . . . . . . . . . . . . . . . . S20

**S6 Parameters Used for Synthetic Data Generation** **S22**

**S7 Additional Parameter Estimates** **S23**
S7.1 Synthetic Data with Two System States . . . . . . . . . . . . . . . . . . S23
S7.2 Synthetic Data with Three System States . . . . . . . . . . . . . . . . . S24
S7.3 Experimental Data: 1 mM . . . . . . . . . . . . . . . . . . . . . . . . . S25
S7.4 Experimental Data: 3 mM . . . . . . . . . . . . . . . . . . . . . . . . . S26
S7.5 Experimental Data: 5 mM . . . . . . . . . . . . . . . . . . . . . . . . . S27
S7.6 Experimental Data: 10 mM . . . . . . . . . . . . . . . . . . . . . . . . S28

**Bibliography** **S29**

### S1 Variables and Notation

In this manuscript, we generally denote *vectors* and *collections* differently, even though mathematically they are very similar objects. Vectors, such as the probability vector $\boldsymbol{\pi}_m$ or the generator matrix $\mathbf{G}$ are bolded. On the other hand, collections, which represent groups of, in some sense, independent objects, such as the trajectory $s_{1:N}$ which is all states $s_n$ grouped together are denoted with the colon notation $i : j$ to denote the range on indices from $i$ to $j$.

| Description | Variable | Units |
|---|---|---|
| The number of pulses | $N$ | - |
| Length of pulse period | $\tau$ | ns |
| The macrotime of the $n$-th pulse | $t_n$ | ns |
| The microtime of the $n$-th pulse | $\mu_n$ | ns or null |
| The measurement of the $n$-th pulse $(\mu_n^d, \mu_n^a)$ | $w_n$ | (ns,ns) |
| Number of states (weak limit in the nonparametric sense) | $M$ | - |
| The $m$-th state | $\sigma_m$ | - |
| The state at the $n$th pulse | $s_n$ | - |
| The system state transition probability matrix | $\mathbf{\Pi}_\sigma$ | - |
| The $m$-th row of $\mathbf{\Pi}_\sigma$ | $\boldsymbol{\pi}_m$ | - |
| Initial state probability vector | $\boldsymbol{\pi}_0$ | - |
| Concentration hyperparameter for $\pi_{1:M}$ and $\pi_0$ | $\alpha$ | - |
| Base distribution over states in the iHMM | $\boldsymbol{\beta}$ | - |
| Concentration hyperparameter for $\beta$ | $\gamma$ | - |
| Donor relaxation rate | $\lambda_d$ | $\text{ns}^{-1}$ |
| Acceptor relaxation rate | $\lambda_a$ | $\text{ns}^{-1}$ |
| FRET rate of $m$-th state | $\lambda_{FRET}^m$ | $\text{ns}^{-1}$ |
| Probability of donor becoming excited by a pulse | $\pi_{ex}$ | - |
| Excitation event at time $n$ | $a_n$ | - |
| Direct acceptor excitation coefficient | $k_a$ | - |
| Efficiency of the donor channel | $\eta_d$ | - |
| Efficiency of the acceptor channel | $\eta_a$ | - |
| Probability of no detector leakage in donor channel | $\phi_{dd}$ | - |
| Probability of no detector leakage in acceptor channel | $\phi_{aa}$ | - |
| Probability of donor channel laser background photon | $p_{bd}$ | - |
| Probability of acceptor channel laser background photon | $p_{ba}$ | - |
| Probability of donor channel uniform background photon | $p_{dd}$ | - |
| Probability of acceptor channel laser background photon | $p_{da}$ | - |
| Donor channel IRF delay mean | $\mu_d^{IRF}$ | ns |
| Donor channel IRF delay variance | $\nu_d^{IRF}$ | $\text{ns}^2$ |
| Acceptor channel IRF delay mean | $\mu_a^{IRF}$ | ns |
| Acceptor channel IRF delay variance | $\nu_a^{IRF}$ | $\text{ns}^2$ |
| The collection of all learned parameters (shorthand) | $\vartheta$ | - |

Table S1: **Table of Variables and Units.** For the convenience of the readers, we include a table with the quantities discussed in this paper and their corresponding symbols.

#### S2 Likelihood for Pulsed Illumination

In order to perform inference over the parameters as described in the main text [1], we use the likelihood described in Eq. 6 of the main text [1]

$$L = p(\boldsymbol{w}|\boldsymbol{\rho}_{start}, \mathbf{\Pi}_\sigma, \mathbf{G}_\psi) \propto \boldsymbol{\rho}_{start}\mathbf{\Pi}_1^\sigma \mathbf{\Pi}_2^\sigma \dots \mathbf{\Pi}_N^\sigma \boldsymbol{\rho}_{norm}^T, \tag{S1}$$

where $\boldsymbol{\rho}_{start}$ is a vector collecting all the initial probabilities. $\mathbf{G}_\psi$ is the photophysical generator matrix given in Eq. 4 of the first companion manuscript [2]. Moreover, $\mathbf{\Pi}_n^\sigma$ is the reduced system state propagator for the $n$-th interpulse period given by

$$\mathbf{\Pi}_n^\sigma = \mathbf{\Pi}_\sigma \odot \mathbf{D}_n^\sigma, \tag{S2}$$

where $\odot$ denotes element-by-element product. Here, $\mathbf{D}_n^\sigma$ is the detection matrix with elements

$$(\mathbf{D}_n^\sigma)_{s_n \to \sigma_j} = p(w_n|s_n, \mathbf{G}_\psi) = \boldsymbol{\rho}_{ground}\mathbf{Q}_n^\psi(s_n)\boldsymbol{\rho}_{norm}^T, \tag{S3}$$

as described in Sec. 3 of the main text [1]. Here, $\boldsymbol{\rho}_{ground}$ is the probability vector where both donor and acceptor are in the ground state. Futhermore, $\mathbf{Q}_n^\psi(s_n)$ is the photophysical propagator for $n$-th interpulse period.

The photophysical propagators take different forms depending on the observation during an interpulse period. To derive the explicit forms of these photophysical propagators, we start from the explicit from of the photophysical generator matrix $\mathbf{G}_\psi$ for a given system state, $s_n$, as

$$\mathbf{G}_\psi = \begin{pmatrix} * & \lambda_{ex}(t) & \lambda_{direct}(t) \\ \lambda_d & * & \lambda_{s_n}^{FRET} \\ \lambda_a & 0 & * \end{pmatrix}, \tag{S4}$$

where $\lambda_{ex}$, $\lambda_{direct}$, $\lambda_d$, $\lambda_a$ and $\lambda_{s_n}^{FRET}$, respectively, denote donor excitation, direct acceptor excitation, donor relaxation, acceptor relaxation and FRET rates. As such, the propagators for empty and nonempty pulses are obtained by replacing the photophysical generator matrix in the generic propagators described in Sec. 2.5.1 of the first companion manuscript [2]

$$\mathbf{Q}_n^\psi = \mathbf{exp}\left(\int_0^{\delta_{pulse}} d\delta \mathbf{G}_\psi^{non}(\delta)\right) \; \mathbf{exp}\Big((\tau - \delta_{pulse})\,\mathbf{G}_\psi^{dark}\Big), \tag{S5}$$

$$\begin{aligned} \mathbf{Q}_n^\psi = \mathbf{exp}\left(\int_0^{\delta_{pulse}} d\delta \mathbf{G}_\psi^{non}(\delta)\right) & \left(\int_0^{\delta_{IRF}} d\epsilon_n \; \mathbf{exp}\left[(\mu_n - \delta_{pulse} - \epsilon_n)\,\mathbf{G}_\psi^{dark}\right] \mathbf{G}_\psi^{rad} \right. \\ & \left. \times \,\mathbf{exp}\left[(\tau - \mu_n + \epsilon_n)\,\mathbf{G}_\psi^{dark}\right] f(\epsilon_n)\right), \end{aligned} \tag{S6}$$

for empty and nonempty pulses, respectively. The different generator matrices above are the reduced forms of $\mathbf{G}^{non}$, $\mathbf{G}^{dark}$ and $\mathbf{G}^{rad}$ introduced in the first companion manuscript [2] Sec. 2.3, now containing only photophysical transitions. In what follows, we will derive these reduced generator matrices and calculate different terms involved in the likelihoods above. We then proceed to take into account the background and instrument response function (IRF) in the likelihoods.

#### S2.1 Excitation

To construct the likelihood for a pulse, we begin by considering the laser pulse itself where we expect no transition other than fluorophore excitation occurring during this period. This assumption is reasonable since pulse duration is too short (often of the order of 100 ps) compared to fluorophore lifetimes. Therefore, the generator matrix for this period is derived from Eq. S4 by setting $\lambda_d = \lambda_a = \lambda_{FRET} = 0$, leading to

$$\mathbf{G}_{\psi}^{non} = \begin{bmatrix} * & \lambda_{ex}(t) & \lambda_{direct}(t) \\ 0 & 0 & 0 \\ 0 & 0 & 0 \end{bmatrix}. \tag{S7}$$

Therefore, the first term in propagators Eq. S5-S6 is obtained as

$$\mathbf{\Pi}_{\psi}^{pulse} = \mathbf{exp}\left( \int_0^{\delta_{pulse}} \begin{bmatrix} * & \lambda_{ex}(t) & \lambda_{direct}(t) \\ 0 & 0 & 0 \\ 0 & 0 & 0 \end{bmatrix} dt \right), \tag{S8}$$

where $\mathbf{\Pi}_{\psi}^{non}$ represents the nonradiative propagator matrix during the laser pulse.

The above expression can be further simplified by taking into account the fact that both excitation rates are proportional to the pulse intensity with different constants of proportionality [3]. Consequently, we can write $\lambda_{direct} = k_a \lambda_{ex}$, where $k_a$ is the ratio of the proportionality constants. The resulting propagator is thus

$$\mathbf{\Pi}_{\psi}^{pulse} = \mathbf{exp}\left( \int_0^{\delta_{pulse}} \begin{bmatrix} * & \lambda_{ex}(t) & k_a\lambda_{ex}(t) \\ 0 & 0 & 0 \\ 0 & 0 & 0 \end{bmatrix} dt \right). \tag{S9}$$

This integral and the subsequent matrix exponential can be solved analytically, with the result

$$\mathbf{\Pi}_{\psi}^{pulse} = \begin{pmatrix} 1 - \pi_{ex} - k_a\pi_{ex} & \pi_{ex} & k_a\pi_{ex} \\ 0 & 0 & 0 \\ 0 & 0 & 0 \end{pmatrix}, \tag{S10}$$

where

$$\pi_{ex} = \frac{1}{1+k_a}\left(1 - \mathbf{exp}\left(-(1+k_a)\int_0^{\delta_{pulse}} \lambda_{ex}(t)dt\right)\right), \tag{S11}$$

where $\pi_{ex}$ and $k_a\pi_{ex}$ are the probabilities that the donor or acceptor is directly excited by the pulse, respectively. This quantity is the same for all the pulses because the molecule is immobilized.

Now, using the obtained propagator for the pulse we can find the photophysical state probability vector immediately after the pulse $\boldsymbol{\rho}_{pulse}$. It is given by

$$\boldsymbol{\rho}_{pulse} = \boldsymbol{\rho}_{ground}\mathbf{\Pi}_{\psi}^{pulse} = \begin{pmatrix} 1 - \pi_{ex} - k_a\pi_{ex}, & \pi_{ex}, & k_a\pi_{ex} \end{pmatrix}, \tag{S12}$$

where $\boldsymbol{\rho}_{ground}$ is the photophysical state probability vector at the beginning of the pulse when both fluorophores are in the ground state by assumption (2) in Sec. 3 of the main text [1].

#### S2.2 Photophysics

Next, we compute the remaining terms in propagators of Eqs. S5-S6. To do so, we first calculate the generator matrices in those terms, namely, $\mathbf{G}_{\psi}^{dark}$ for no photon detection, and $\mathbf{G}_{\psi}^{rad}$ for photon detection. These two generators describe events after the laser pulse and before the next laser pulse where no fluorophore excitation may take place and thus we have $\lambda_{ex} = \lambda_{direct} = 0$.

Now, for an empty interpulse period where there is no photon detection, there is still a chance for emitted photons that are not detected quantified by detector efficiencies $\eta_d$ and $\eta_a$ for donor and acceptor channels, respectively. Therefore, we can write (see Sec. 2.5.1 in the first companion manuscript [2])

$$\mathbf{G}_{\psi}^{dark} = \begin{pmatrix} 0 & 0 & 0 \\ (1-\eta_d)\lambda_d & -\lambda_d - \lambda_{s_n}^{FRET} & \lambda_{s_n}^{FRET} \\ (1-\eta_a)\lambda_a & 0 & -\lambda_a \end{pmatrix}. \tag{S13}$$

For nonempty interpulse periods, only radiative transitions associated with the detected photon are possible at that detection moment, therefore we further set the nonradiative transition rates $\lambda_{FRET} = 0$. If a photon is detected in the donor channel, the radiative propagator is thus

$$\mathbf{G}_{\psi}^{rad(D)} = \begin{pmatrix} 0 & 0 & 0 \\ \eta_d\phi_{dd}\lambda_d & 0 & 0 \\ \eta_d(1-\phi_{aa})\lambda_a & 0 & 0 \end{pmatrix}, \tag{S14}$$

and for the acceptor channel

$$\mathbf{G}_{\psi}^{rad(A)} = \begin{pmatrix} 0 & 0 & 0 \\ \eta_a(1-\phi_{dd})\lambda_d & 0 & 0 \\ \eta_a\phi_{aa}\lambda_a & 0 & 0 \end{pmatrix}, \tag{S15}$$

where $(1-\phi_{dd})$ and $(1-\phi_{aa})$ denote the crosstalk probabilities for donor and acceptor channels, respectively.

Now, if we ignore the background and the IRF for the moment, using the obtained generators above, the elements of the detection matrix $(\mathbf{D}_n^{\sigma})_{s_n \to \sigma_j}$ for an empty pulse, a nonempty pulse with a donor photon, and a nonempty pulses with an acceptor photon are, respectively, given as

$$(\mathbf{D}_n^{\sigma})_{s_n \to \sigma_j}(\emptyset, \emptyset) = \boldsymbol{\rho}_{pulse}\, \mathbf{exp}(\tau \mathbf{G}_{\psi}^{dark}) \boldsymbol{\rho}_{norm}^T, \quad \text{no photon,} \tag{S16}$$

$$(\mathbf{D}_n^{\sigma})_{s_n \to \sigma_j}(\mu, \emptyset) = \boldsymbol{\rho}_{pulse}\, \mathbf{exp}(\mu_n \mathbf{G}_{\psi}^{dark}) \mathbf{G}_{\psi}^{rad(D)}\, \mathbf{exp}((\tau - \mu_n)\, \mathbf{G}_{\psi}^{dark}) \boldsymbol{\rho}_{norm}^T, \quad \text{donor photon} \tag{S17}$$

$$(\mathbf{D}_n^{\sigma})_{s_n \to \sigma_j}(\emptyset, \mu) = \boldsymbol{\rho}_{pulse}\, \mathbf{exp}(\mu_n \mathbf{G}_{\psi}^{dark}) \mathbf{G}_{\psi}^{rad(A)}\, \mathbf{exp}((\tau - \mu_n)\, \mathbf{G}_{\psi}^{dark}) \boldsymbol{\rho}_{norm}^T, \quad \text{acceptor photon} \tag{S18}$$

where we ignored the integrals due to IRF in Eq. S5-S6. Moreover, $\emptyset$ and $\mu$ as the first input, respectively denote no photon and a photon with arrival time $\mu$ from the donor channel. The same applies to the second input but for the acceptor channel. These elements can be analytically solved as

$$
(\mathbf{D}_n^{\sigma})_{s_n\to\sigma_j}(\emptyset,\emptyset) = \\
\boldsymbol{\rho}_{pulse}\begin{pmatrix} 1 & 0 & 0 \\ A(\tau) & \mathbf{exp}(-(\lambda_d+\lambda_{s_n}^{FRET})\tau) & B(\tau) \\ (1-\eta_a)(1-\mathbf{exp}(-\lambda_a\tau)) & 0 & \mathbf{exp}(-\lambda_a\tau)\end{pmatrix}\boldsymbol{\rho}_{norm}^T, \tag{S19}
$$

$$
(\mathbf{D}_n^{\sigma})_{s_n\to\sigma_j}(\mu,\emptyset) = \\
\boldsymbol{\rho}_{pulse}\begin{pmatrix} 0 & 0 & 0 \\ \eta_d\phi_{dd}\lambda_d\,\mathbf{exp}(-(\lambda_d+\lambda_{s_n}^{FRET})\mu)+\eta_d(1-\phi_{aa})\lambda_a B(\mu) & 0 & 0 \\ \eta_d(1-\phi_{aa})\lambda_a\,\mathbf{exp}(-\lambda_a\mu) & 0 & 0\end{pmatrix}\boldsymbol{\rho}_{norm}^T, \tag{S20}
$$

$$
(\mathbf{D}_n^{\sigma})_{s_n\to\sigma_j}(\emptyset,\mu) = \\
\boldsymbol{\rho}_{pulse}\begin{pmatrix} 0 & 0 & 0 \\ \eta_a(1-\phi_{dd})\lambda_d\,\mathbf{exp}(-(\lambda_d+\lambda_{s_n}^{FRET})\mu)+\eta_a\phi_{aa}\lambda_a B(\mu) & 0 & 0 \\ \eta_a\phi_{aa}\lambda_a\,\mathbf{exp}(-\lambda_a\mu) & 0 & 0\end{pmatrix}\boldsymbol{\rho}_{norm}^T, \tag{S21}
$$

where $\tau$ is the interpulse period and

$$
A(t) = \frac{(1-\eta_d)\lambda_d+(1-\eta_a)\lambda_{s_n}^{FRET}}{\lambda_d+\lambda_{s_n}^{FRET}}(1-\mathbf{exp}(-(\lambda_d+\lambda_{s_n}^{FRET})t)), \tag{S22}
$$

$$
B(t) = \frac{\lambda_{s_n}^{FRET}}{-\lambda_d-\lambda_{s_n}^{FRET}+\lambda_a}(\mathbf{exp}(-(\lambda_d+\lambda_{s_n}^{FRET})t)-\mathbf{exp}(-\lambda_a t)). \tag{S23}
$$

These can be further simplified by making the assumption that interpulses period is long in comparison to the fluorophore lifetimes (assumption (2) above). In essence, we take $\tau\to\infty$. Therefore, the elements of the detection matrix when no photon is detected becomes

$$
\lim_{\tau\to\infty}(\mathbf{D}_n^{\sigma})_{s_n\to\sigma_j}(\emptyset,\emptyset) = \boldsymbol{\rho}_{pulse}\begin{pmatrix} 1 & 0 & 0 \\ \frac{(1-\eta_d)\lambda_d+(1-\eta_a)\lambda_{s_n}^{FRET}}{\lambda_d+\lambda_{s_n}^{FRET}} & 0 & 0 \\ (1-\eta_a) & 0 & 0\end{pmatrix}\boldsymbol{\rho}_{norm}^T. \tag{S24}
$$

Now, since $\boldsymbol{\rho}_{norm} = [1,\,1,\,1]$, these matrices can be reduced to vectors by incorporating $\boldsymbol{\rho}_{norm}^T$ as

$$
(\mathbf{D}_n^{\sigma})_{s_n\to\sigma_j}(\emptyset,\emptyset) = \boldsymbol{\rho}_{pulse}\begin{pmatrix} 1 \\ \frac{(1-\eta_d)\lambda_d+(1-\eta_a)\lambda_{s_n}^{FRET}}{\lambda_d+\lambda_{s_n}^{FRET}} \\ (1-\eta_a)\end{pmatrix}, \tag{S25}
$$

$$
(\mathbf{D}_n^{\sigma})_{s_n\to\sigma_j}(\mu,\emptyset) = \boldsymbol{\rho}_{pulse}\begin{pmatrix} 0 \\ \eta_d\phi_{dd}\lambda_d\,\mathbf{exp}(-(\lambda_d+\lambda_{s_n}^{FRET})\mu)+\eta_d(1-\phi_{aa})\lambda_a B(\mu) \\ \eta_d(1-\phi_{aa})\lambda_a\,\mathbf{exp}(-\lambda_a\mu)\end{pmatrix}, \tag{S26}
$$

$$
(\mathbf{D}_n^{\sigma})_{s_n\to\sigma_j}(\emptyset,\mu) = \boldsymbol{\rho}_{pulse}\begin{pmatrix} 0 \\ \eta_a(1-\phi_{dd})\lambda_d\,\mathbf{exp}(-(\lambda_d+\lambda_{s_n}^{FRET})\mu)+\eta_a\phi_{aa}\lambda_a B(\mu) \\ \eta_a\phi_{aa}\lambda_a\,\mathbf{exp}(-\lambda_a\mu)\end{pmatrix}. \tag{S27}
$$

Additionally, it is sometimes convenient to consider the likelihood of only detecting a donor or acceptor photon, regardless of the photon arrival time. We find this by marginalizing over the arrival times, and denote these marginalized likelihoods by

$$(\hat{\mathbf{D}}_n^\sigma)_{s_n\to\sigma_j}(d) = \int_0^\infty (\mathbf{D}_n^\sigma)_{s_n\to\sigma_j}(t,\emptyset)dt = \boldsymbol{\rho}_{pulse}\begin{pmatrix} 0 \\ \eta_d\phi_{dd}(1-\varepsilon_{s_n}^{FRET}) + \eta_d(1-\phi_{aa})\varepsilon_{s_n}^{FRET} \\ \eta_d(1-\phi_{aa}) \end{pmatrix}, \tag{S28}$$

$$(\hat{\mathbf{D}}_n^\sigma)_{s_n\to\sigma_j}(a) = \int_0^\infty (\mathbf{D}_n^\sigma)_{s_n\to\sigma_j}(\emptyset,t)dt = \boldsymbol{\rho}_{pulse}\begin{pmatrix} 0 \\ \eta_d(1-\phi_{dd})(1-\varepsilon_{s_n}^{FRET}) + \eta_d\phi_{aa}\varepsilon_{s_n}^{FRET} \\ \eta_a\phi_{aa} \end{pmatrix}, \tag{S29}$$

where $\varepsilon_{s_n}^{FRET} = \lambda_{s_n}^{FRET}/(\lambda_d + \lambda_{FRET}^{s_n})$ is the FRET efficiency for the system state $s_n$. Here, $\hat{\mathbf{D}}_n^\sigma$ denotes marginalization over arrival times.

In what follows, we will describe how to include the IRF and background into the derived detection matrices in this section.

#### S2.3 Instrument Response Function

The IRF refers to the delay between a photon arrival to a detector and the arrival time reported by the detector due to the electronics. We incorporate it by concluding that the reported arrival time $t_{rep}$ is the sum of two random variables, $t_{arrive}$ and $t_{IRF}$, as follows

$$t_{rep} = t_{arrive} + t_{IRF}. \tag{S30}$$

As it is a sum of two random variables the resulting distribution of $t_{rep}$ is a convolution of the photon arrival time distribution with the IRF distribution. Here, we assume that the IRF is distributed according to

$$t_{IRF} \sim \mathbf{Normal}(\mu_{IRF}, \nu_{IRF}), \tag{S31}$$

with each channel having a unique mean $\mu_{IRF}$ and variance $\nu_{IRF}$. Moreover, the distribution of $t_{arrive}$ is described by $(\mathbf{D}_n^\sigma)_{s_n\to\sigma_j}$ derived in the previous section.

Now, we can obtain the likelihood in the presence of the IRF by calculating the convolution implied by Eq. S30. That is obtained as follows

$$(\mathbf{D}_n^\sigma)_{s_n\to\sigma_j}^{IRF}(\mu,\emptyset) = \boldsymbol{\rho}_{pulse}\begin{pmatrix} 0 \\ \eta_d\phi_{dd}\lambda_d f_d(\mu,\lambda_d+\lambda_{s_n}^{FRET}) + \eta_d(1-\phi_{aa})\lambda_a B_{f_d}(\mu) \\ \eta_d(1-\phi_{aa})\lambda_a f_d(\mu,\lambda_a) \end{pmatrix}, \tag{S32}$$

$$(\mathbf{D}_n^\sigma)_{s_n\to\sigma_j}^{IRF}(\emptyset,\mu) = \boldsymbol{\rho}_{pulse}\begin{pmatrix} 0 \\ \eta_a(1-\phi_{dd})\lambda_d f_a(\mu,\lambda_d+\lambda_{s_n}^{FRET}) + \eta_a\phi_{aa}\lambda_a B_{f_a}(\mu) \\ \eta_a\phi_{aa}\lambda_a f_a(\mu,\lambda_a) \end{pmatrix}, \tag{S33}$$

where

$$B_{f_d}(t) = \frac{\lambda_{s_n}^{FRET}}{-\lambda_d - \lambda_{s_n}^{FRET} + \lambda_a}(f_d(t, \lambda_d + \lambda_{s_n}^{FRET}) - f_d(t, \lambda_a)), \tag{S34}$$

$$f_d(t, \lambda) = \frac{1}{2}\mathbf{exp}\left(\frac{\lambda}{2}(2\mu_d^{IRF} + \lambda\nu_d^{IRF} - 2t)\right)\mathbf{erfc}\left(\frac{\mu_d^{IRF} + \lambda\nu_d^{IRF} - t}{\sqrt{2\nu_d^{IRF}}}\right), \tag{S35}$$

$$B_{f_a}(t) = \frac{\lambda_{s_n}^{FRET}}{-\lambda_d - \lambda_{s_n}^{FRET} + \lambda_a}(f_a(t, \lambda_d + \lambda_{s_n}^{FRET}) - f_a(t, \lambda_a)), \tag{S36}$$

$$f_a(t, \lambda) = \frac{1}{2}\mathbf{exp}\left(\frac{\lambda}{2}(2\mu_a^{IRF} + \lambda\nu_a^{IRF} - 2t)\right)\mathbf{erfc}\left(\frac{\mu_a^{IRF} + \lambda\nu_a^{IRF} - t}{\sqrt{2\nu_a^{IRF}}}\right), \tag{S37}$$

where $\mathbf{erfc}(\cdot) = 1 - \mathbf{erf}(\cdot)$ is the *complementary error function*. Moreover, note that $(\mathbf{D}_n^{\sigma})_{s_n\to\sigma_j}^{IRF}(\emptyset, \emptyset) = (\mathbf{D}_n^{\sigma})_{s_n\to\sigma_j}(\emptyset, \emptyset)$ since there is no photon and thus no IRF effect.

#### S2.4 Background

In this section, we proceed to include background emissions in our formulation following Sec. 2.6 of the first comapnion manuscript [2]. The background photons come from extra light sources present in the environment in addition to the FRET pair. Such source of photon is, in general, characterized by two components: 1) photon emission probabilities for each channel; and 2) distribution of photon arrival times over the interpulse window.

Here, we first assume $p_d$ and $p_a$ to be the probability that a photon is emitted in the donor and acceptor channels, respectively. Further, let $g_d(t)$ and $g_a(t)$ be probability density functions that describe the distribution of background photons' arrival times within the interpulse window for the donor and acceptor channels, respectively. Moreover, note that if the source is such that there is some relationship between donor and acceptor photons, we would additionally require a joint probability distribution $g_{da}(t_d, t_a)$, but in the case of background, we assume that the channels are independent. Therefore, the distribution over measurements for this source is

$$p_{bg}(w_n) = \begin{cases} (1-p_d)(1-p_a) & w_n = (\emptyset, \emptyset) \\ p_d g_d(\mu_d)(1-p_a) & w_n = (\mu_d, \emptyset) \\ (1-p_d)p_a g_a(\mu_a) & w_n = (\emptyset, \mu_a) \\ p_{bd}p_{ba}g_d(\mu_d)g_a(\mu_a) & w_n = (\mu_d, \mu_a). \end{cases} \tag{S38}$$

In the presence of a background source, we run into the complication that, in most single photon pulsed illumination setups, only the first photon arriving to a detector channel is recorded. This means that there is a competition between photons from different sources, namely, donor fluorophore, acceptor fluorophore, and background, to first reaching the detector. In many cases, this effect can be ignored, but here we take it into account for generality. In this case, we can write the likelihood in the presence of background but absence of the

IRF, $(\mathbf{D}_n^\sigma)^{bg}_{s_n\to\sigma_j}$, as follows

$$(\mathbf{D}_n^\sigma)^{bg}_{s_n\to\sigma_j}(w_n) =$$
$$\begin{cases} p_{bg}(\emptyset,\emptyset)(\mathbf{D}_n^\sigma)_{s_n\to\sigma_j}(\emptyset,\emptyset) & w_n=(\emptyset,\emptyset)\\ p_{bg}(\emptyset,\emptyset)(\mathbf{D}_n^\sigma)_{s_n\to\sigma_j}(\mu_d,\emptyset)+p_{bg}(\mu_d,\emptyset)(\mathbf{D}_n^\sigma)_{s_n\to\sigma_j}(\emptyset,\emptyset)+m_d(\mu_d,\emptyset) & w_n=(\mu_d,\emptyset)\\ p_{bg}(\emptyset,\emptyset)(\mathbf{D}_n^\sigma)_{s_n\to\sigma_j}(\emptyset,\mu_a)+p_{bg}(\emptyset,\mu_a)(\mathbf{D}_n^\sigma)_{s_n\to\sigma_j}(\emptyset,\emptyset)+m_a(\emptyset,\mu_a) & w_n=(\emptyset,\mu_a)\\ p_{bg}(\emptyset,\mu_a)(\mathbf{D}_n^\sigma)_{s_n\to\sigma_j}(\mu_d,\emptyset)+p_{bg}(\mu_d,\emptyset)(\mathbf{D}_n^\sigma)_{s_n\to\sigma_j}(\emptyset,\mu_a) & w_n=(\mu_d,\mu_a)\\ +p_{bg}(\mu_d,\mu_a)(\mathbf{D}_n^\sigma)_{s_n\to\sigma_j}(\emptyset,\emptyset)+M_d(\mu_d,\mu_a)+M_a(\mu_d,\mu_a), & \end{cases} \tag{S39}$$

where $p_{bg}$ is given by Eq. S38 and $(\mathbf{D}_n^\sigma)_{s_n\to\sigma_j}$ is the likelihood for signal photons, *i.e.*, photons from fluorophores. Further, $m_d$, $m_a$, $M_d$ and $M_a$ correspond to the cases where both background and signal photon are present, but only the smaller arrival time is detected. These are derived by finding the distribution of the minimum arrival times between the competing photons as follows

$$\begin{aligned} m_d(t,\emptyset) =& p_d(1-p_a)\Bigg[(g_d(t)(\hat{\mathbf{D}}_n^\sigma)_{s_n\to\sigma_j}(d)+(\mathbf{D}_n^\sigma)_{s_n\to\sigma_j}(t,\emptyset)-g_d(t)\left(\int_0^t(\mathbf{D}_n^\sigma)_{s_n\to\sigma_j}(s,\emptyset)ds\right)\\ &-\left(\int_0^t g_d(s)ds\right)(\mathbf{D}_n^\sigma)_{s_n\to\sigma_j}(t,\emptyset)\Bigg], \end{aligned} \tag{S40}$$

$$\begin{aligned} m_a(\emptyset,t) =& (1-p_d)p_a\Bigg[g_a(t)(\hat{\mathbf{D}}_n^\sigma)_{s_n\to\sigma_j}(a)+(\mathbf{D}_n^\sigma)_{s_n\to\sigma_j}(t,\emptyset)-g_a(t)\left(\int_0^t(\mathbf{D}_n^\sigma)_{s_n\to\sigma_j}(s,\emptyset)ds\right)\\ &-\left(\int_0^t g_a(s)ds\right)(\mathbf{D}_n^\sigma)_{s_n\to\sigma_j}(t,\emptyset)\Bigg], \end{aligned} \tag{S41}$$

$$\begin{aligned} M_d(t_d,t_a) =& p_dp_a\Bigg[g_d(t_d)(\hat{\mathbf{D}}_n^\sigma)_{s_n\to\sigma_j}(d)+(\mathbf{D}_n^\sigma)_{s_n\to\sigma_j}(t_d,\emptyset)-g_d(t)\left(\int_0^{t_d}(\mathbf{D}_n^\sigma)_{s_n\to\sigma_j}(s,\emptyset)ds\right)\\ &-\left(\int_0^{t_d} g_d(s)ds\right)(\mathbf{D}_n^\sigma)_{s_n\to\sigma_j}(t_d,\emptyset)\Bigg]g_a(t_a), \end{aligned} \tag{S42}$$

$$\begin{aligned} M_a(t_d,t_a) =& p_dp_ag_d(t_d)\Bigg[g_a(t_a)(\hat{\mathbf{D}}_n^\sigma)_{s_n\to\sigma_j}(a)+(\mathbf{D}_n^\sigma)_{s_n\to\sigma_j}(t_a,\emptyset)-g_a(t_a)\left(\int_0^{t_a}(\mathbf{D}_n^\sigma)_{s_n\to\sigma_j}(s,\emptyset)ds\right)\\ &-\left(\int_0^{t_a} g_a(s)ds\right)(\mathbf{D}_n^\sigma)_{s_n\to\sigma_j}(t_a,\emptyset)\Bigg], \end{aligned} \tag{S43}$$

where $(\hat{\mathbf{D}}_n^\sigma)_{s_n\to\sigma_j}$ is the marginalized element introduced in Eq. S28.

Using this general method of incorporating additional light sources into our framework, we can add the two most prominant background sources observed in the data: 1) laser photons which are distributed the same as laser pulse and termed laser background; and 2) uniform background which are uniformly distributed over the interpulse window and termed uniform background. In what follows, we will discuss the inclusion of these two backgrounds in our model.

###### S2.4.1 Laser background

The primary goal of this section is constructing a general method for adding background photons originating from the laser source to our pulsed illumination framework. These photons arrive to the detector distributed according to the intensity of the laser pulse across the interpulse window. Moreover, since the pulse width is extremely narrow, it can be effectively considered as a delta function. Therefore, using our description for a generic background source (Eq. S38), we can describe this laser background as

$$p_{bg}(w_n) = \begin{cases} (1-p_{bd})(1-p_{ba}) & w_n = (\emptyset, \emptyset) \\ p_{bd}\delta(\mu_d)(1-p_{ba}) & w_n = (\mu_d, \emptyset) \\ (1-p_{bd})p_{ba}\delta(\mu_a) & w_n = (\emptyset, \mu_a) \\ p_{bd}p_{ba}\delta(\mu_d)\delta(\mu_a) & w_n = (\mu_d, \mu_a) \end{cases}, \tag{S44}$$

where we used $g_{d\backslash a}(\mu_{d\backslash a}) = \delta(\mu_{d\backslash a})$ for laser photons.

Since the laser photons arrive exactly at the beginning of the interpulse window, they are going to naturally win the competition between multiple present photons from different sources. This in turn simplifies the terms $m_d$, $m_a$, $M_d$, and $M_a$ in Eq. S40-S43 for laser photons as follows

$$m^l_d(\mu_d, \emptyset) = p_{bd}(1-p_{ba})(\hat{\mathbf{D}}^\sigma_n)_{s_n\to\sigma_j}(d)\delta(\mu_d), \tag{S45}$$

$$m^l_a(\emptyset, \mu_a) = (1-p_{bd})p_{ba}(\hat{\mathbf{D}}^\sigma_n)_{s_n\to\sigma_j}(a)\delta(\mu_a), \tag{S46}$$

$$M^l_d(\mu_d, \mu_a) = p_{bd}p_{ba}(\hat{\mathbf{D}}^\sigma_n)_{s_n\to\sigma_j}(d)\delta(\mu_d)\delta(\mu_a), \tag{S47}$$

$$M^l_a(\mu_d, \mu_a) = p_{bd}p_{ba}(\hat{\mathbf{D}}^\sigma_n)_{s_n\to\sigma_j}(a)\delta(\mu_d)\delta(\mu_a). \tag{S48}$$

Here, the marginalized terms $(\hat{\mathbf{D}}^\sigma_n)_{s_n\to\sigma_j}$ (defined in Eq. S28) account for the probability of receiving a signal photon from the fluorophore, even if it is not detected due to the laser background photon arriving first.

Now, by substituting Eq. S44-S48 in Eq. S39 we can derive the likelihood model of the photons reaching to the detector in the presence of laser photons. To derive the reported arrival time likelihood model, we still need to add the IRF effect. To do so, we need to convolve the IRF with the delta function that describes the laser photon distributions across the interpulse window. This results in the IRF itself which is given by a Normal distribution

$$\int d\omega \delta(t-\omega)\mathbf{Normal}(\omega; \mu_{IRF}, \nu_{IRF}) = \mathbf{Normal}(t; \mu_{IRF}, \nu_{IRF}), \tag{S49}$$

where $\omega$ is an auxiliary variable. As such, using the $(\mathbf{D}^\sigma_n)^{IRF}_{s_n\to\sigma_j}$ in Eqs. S32-S33 and the background terms as described above, we obtain the likelihood model in the presence of laser background $(\mathbf{D}_n^\sigma)_{s_n\to\sigma_j}^{laser}$ as

$$(\mathbf{D}_n^\sigma)_{s_n\to\sigma_j}^{laser}(\emptyset,\emptyset) = (1-p_{bd})(1-p_{ba})(\mathbf{D}_n^\sigma)_{s_n\to\sigma_j}(\emptyset,\emptyset), \tag{S50}$$

$$\begin{aligned}(\mathbf{D}_n^\sigma)_{s_n\to\sigma_j}^{laser}(\mu_d,\emptyset) =&(1-p_{bd})(1-p_{ba})(\mathbf{D}_n^\sigma)_{s_n\to\sigma_j}^{IRF}(\mu_d,\emptyset)\\ &+ p_{bd}\mathbf{Normal}(\mu_d;\mu_d^{IRF},\nu_d^{IRF})(1-p_{ba})(\mathbf{D}_n^\sigma)_{s_n\to\sigma_j}(\emptyset,\emptyset)\\ &+ p_{bd}(1-p_{ba})(\hat{\mathbf{D}}_n^\sigma)_{s_n\to\sigma_j}(d)\mathbf{Normal}(\mu_d;\mu_d^{IRF},\nu_d^{IRF}),\end{aligned} \tag{S51}$$

$$\begin{aligned}(\mathbf{D}_n^\sigma)_{s_n\to\sigma_j}^{laser}(\emptyset,\mu_a) =&(1-p_{bd})(1-p_{ba})(\mathbf{D}_n^\sigma)_{s_n\to\sigma_j}^{IRF}(\emptyset,\mu_a)\\ &+ (1-p_{bd})p_{ba}\mathbf{Normal}(\mu_a;\mu_a^{IRF},\nu_a^{IRF})(\mathbf{D}_n^\sigma)_{s_n\to\sigma_j}(\emptyset,\emptyset)\\ &+ (1-p_{bd})p_{ba}(\hat{\mathbf{D}}_n^\sigma)_{s_n\to\sigma_j}(a)\mathbf{Normal}(\mu_a;\mu_a^{IRF},\nu_a^{IRF}),\end{aligned} \tag{S52}$$

$$\begin{aligned}(\mathbf{D}_n^\sigma)_{s_n\to\sigma_j}^{laser}(\mu_d,\mu_a) =&(1-p_{bd})p_{ba}\mathbf{Normal}(\mu_a;\mu_a^{IRF},\nu_a^{IRF})(\mathbf{D}_n^\sigma)_{s_n\to\sigma_j}^{IRF}(\mu_d,\emptyset)\\ &+ p_{bd}\mathbf{Normal}(\mu_d;\mu_d^{IRF},\nu_d^{IRF})(1-p_{ba})(\mathbf{D}_n^\sigma)_{s_n\to\sigma_j}^{IRF}(\emptyset,\mu_a)\\ &+ p_{bd}p_{ba}\mathbf{Normal}(\mu_d;\mu_d^{IRF},\nu_d^{IRF})\mathbf{Normal}(\mu_a;\mu_a^{IRF},\nu_a^{IRF})\\ &\times((\hat{\mathbf{D}}_n^\sigma)_{s_n\to\sigma_j}(d)+(\hat{\mathbf{D}}_n^\sigma)_{s_n\to\sigma_j}(a)).\end{aligned} \tag{S53}$$

##### S2.4.2 Uniform background

Finally, we incorporate uniform background, which represents the combination of all ambient light sources that emit photons with a constant rate, independent of the laser pulses. We introduce uniform background after the IRF as the arrival time distribution of these photons is not affected by the IRF, remaining uniform over the entire interpulse window. Once again, using the form for a generic light source from Eq. S38, we describe uniform background as

$$p_{dbg}(w_n) = \begin{cases}(1-p_{dd})(1-p_{da}) & w_n=(\emptyset,\emptyset)\\ p_{dd}(1/\tau)(1-p_{da}) & w_n=(\mu_d,\emptyset)\\ (1-p_{dd})p_{da}(1/\tau) & w_n=(\emptyset,\mu_a)\\ p_{dd}p_{da}(1/\tau)^2 & w_n=(\mu_d,\mu_a)\end{cases}. \tag{S54}$$

Combining this source with $(\mathbf{D}_n^\sigma)_{s_n\to\sigma_j}^{laser}$ from the previous section in the same way as described in Eq. S39, we arrive at our final expression for the detection matrices as

$$(\mathbf{D}_n^\sigma)_{s_n\to\sigma_j}(w_n) = \begin{cases} p_{dbg}(\emptyset,\emptyset)(\mathbf{D}_n^\sigma)_{s_n\to\sigma_j}^{laser}(\emptyset,\emptyset) & w_n=(\emptyset,\emptyset)\\ p_{dbg}(\emptyset,\emptyset)(\mathbf{D}_n^\sigma)_{s_n\to\sigma_j}^{laser}(\mu_d,\emptyset)+p_{dbg}(\mu_d,\emptyset)(\mathbf{D}_n^\sigma)_{s_n\to\sigma_j}^{laser}(\emptyset,\emptyset)+m_d^u(\mu_d,\emptyset) & w_n=(\mu_d,\emptyset)\\ p_{dbg}(\emptyset,\emptyset)(\mathbf{D}_n^\sigma)_{s_n\to\sigma_j}^{laser}(\emptyset,\mu_a)+p_{dbg}(\emptyset,\mu_a)(\mathbf{D}_n^\sigma)_{s_n\to\sigma_j}^{laser}(\emptyset,\emptyset)+m_a^u(\emptyset,\mu_a) & w_n=(\emptyset,\mu_a)\\ p_{dbg}(\emptyset,\mu_a)(\mathbf{D}_n^\sigma)_{s_n\to\sigma_j}^{laser}(\mu_d,\emptyset)+p_{dbg}(\mu_d,\emptyset)(\mathbf{D}_n^\sigma)_{s_n\to\sigma_j}^{laser}(\emptyset,\mu_a) & w_n=(\mu_d,\mu_a)\\ +p_{dbg}(\mu_d,\mu_a)(\mathbf{D}_n^\sigma)_{s_n\to\sigma_j}^{laser}(\emptyset,\emptyset)+M_d^u(\mu_d,\mu_a)+M_a^u(\mu_d,\mu_a). & \end{cases} \tag{S55}$$

Unlike for the laser background, the terms $m^u_d$, $m^u_a$, $M^u_d$, and $M^u_a$ are no longer simple to compute. Therefore, here, we present their approximate form

$$m^u_d(t,\emptyset) \approx p_{dd}(1-p_{da})\left[\frac{1}{\tau}(\hat{\mathbf{D}}^\sigma_n)^{laser}_{s_n\to\sigma_j}(d) - \frac{1}{\tau}\left(\int_0^t (\mathbf{D}^\sigma_n)^{laser}_{s_n\to\sigma_j}(s,\emptyset)ds\right) \right.$$
$$\left. + \left(1-\frac{t}{\tau}\right)(\mathbf{D}^\sigma_n)^{laser}_{s_n\to\sigma_j}(t,\emptyset)\right], \tag{S56}$$

$$m^u_a(\emptyset,t) \approx (1-p_{dd})p_{da}\left[\frac{1}{\tau}(\hat{\mathbf{D}}^\sigma_n)^{laser}_{s_n\to\sigma_j}(a) - \frac{1}{\tau}\left(\int_0^t (\mathbf{D}^\sigma_n)^{laser}_{s_n\to\sigma_j}(s,\emptyset)ds\right) \right.$$
$$\left. + \left(1-\frac{t}{\tau}\right)(\mathbf{D}^\sigma_n)^{laser}_{s_n\to\sigma_j}(t,\emptyset)\right], \tag{S57}$$

$$M^u_d(t_d,t_a) \approx p_{dd}p_{da}\left[\frac{1}{\tau}(\hat{\mathbf{D}}^\sigma_n)^{laser}_{s_n\to\sigma_j}(d) - \frac{1}{\tau}\left(\int_0^{t_d} (\mathbf{D}^\sigma_n)^{laser}_{s_n\to\sigma_j}(s,\emptyset)ds\right) \right.$$
$$\left. + \left(1-\frac{t}{\tau}\right)(\mathbf{D}^\sigma_n)^{laser}_{s_n\to\sigma_j}(t_d,\emptyset)\right]\frac{1}{\tau}, \tag{S58}$$

$$M^u_a(t_d,t_a) \approx p_{dd}p_{da}\frac{1}{\tau}\left[\frac{1}{\tau}(\hat{\mathbf{D}}^\sigma_n)^{laser}_{s_n\to\sigma_j}(a) - \frac{1}{\tau}\left(\int_0^{t_a} (\mathbf{D}^\sigma_n)^{laser}_{s_n\to\sigma_j}(s,\emptyset)ds\right) \right.$$
$$\left. + \left(1-\frac{t}{\tau}\right)(\mathbf{D}^\sigma_n)^{laser}_{s_n\to\sigma_j}(t_a,\emptyset)\right]. \tag{S59}$$

Here, we derived the most general likelihood for smFRET under pulsed illumination and used it in our analysis. However, as mentioned earlier, this likelihood can be much simplified to an approximate form by ignoring the terms associated to the competitions between photons reaching to the detectors in $(\mathbf{D}^\sigma_n)^{laser}_{s_n\to\sigma_j}$ (Eqs. S50-S53) and $(\mathbf{D}^\sigma_n)_{s_n\to\sigma_j}$ (Eq. S59).

### S3 Model Structure and Priors

After deriving the likelihood, in this section, we present our priors to derive the parametric and nonparametric posteriors. The parameters are arranged according to their hierarchical dependency, meaning that if a parameter depends on another, it will necessarily come after it. Parameters whose priors' parameters are set by hand rather than by another parameter are highlighted with a $(*)$.

#### S3.1 Parametric Model

$$
\begin{aligned}
\boldsymbol{\pi}_0 &\sim \mathbf{Dirichlet}(1, 1, \ldots, 1), && (*)\\
\boldsymbol{\pi}_m &\sim \mathbf{Dirichlet}(1, 1, \ldots, 1), && m \in \{1, \ldots, M\},\ (*)\\
s_1 &\sim \mathbf{Categorical}(\pi_0), &&\\
s_n|s_{n-1} &\sim \mathbf{Categorical}(\pi_{s_{n-1}}), && n \in \{2, 3, \ldots, N\},\\
\pi_{ex} &\sim \mathbf{Beta}(1, 1), && (*)\\
a_n &\sim \mathbf{Categorical}(1 - \pi_{ex} - k_a\pi_{ex},\ \pi_{ex},\ k_a\pi_{ex}), && n \in \{1, \ldots, N\},\\
\lambda_d &\sim \mathbf{Gamma}(1, 1), && (*)\\
\lambda_a &\sim \mathbf{Gamma}(1, 1), && (*)\\
\lambda_{\sigma_m}^{FRET} &\sim \mathbf{Gamma}(1, 1), && m \in \{1, \ldots, M\},\ (*)\\
w_n &\sim p(w|a_n, \lambda_d, \lambda_a, \lambda_{s_n}^{FRET}), && n \in \{1, \ldots, N\},
\end{aligned}
$$

where the distribution $p(w|a_n, \lambda_d, \lambda_a, \lambda_{s_n}^{FRET})$ is the likelihood derived in Section S2 with $\boldsymbol{\rho}_{pulse}$ set by the auxiliary parameter $a_n$.

#### S3.2 Nonparametric Model

$$
\begin{aligned}
\gamma &\sim \textbf{Gamma}(1,1), && (*)\\
\beta &\sim \textbf{Dirichlet}\left(\frac{\gamma}{M},\dots,\frac{\gamma}{M}\right),\\
\alpha &\sim \textbf{Gamma}(1,1), && (*)\\
\kappa &\sim \textbf{Beta}(\phi,1), && (*)\\
\boldsymbol{\pi}_0 &\sim \textbf{Dirichlet}(\alpha\boldsymbol{\beta}),\\
\boldsymbol{d}_m &= \begin{cases}(d_m)_i = 1 & i = m\\ (d_m)_i = 0 & i \neq m\end{cases},\\
\boldsymbol{\pi}_m &\sim \textbf{Dirichlet}(\alpha((1-\kappa)\boldsymbol{\beta}+\kappa\boldsymbol{d}_m)), && m\in\{1,\dots,M\},\\
s_1 &\sim \textbf{Categorical}(\boldsymbol{\pi}_0),\\
s_n|s_{n-1} &\sim \textbf{Categorical}(\boldsymbol{\pi}_{s_{n-1}}), && n\in\{2,3,\dots,N\},\\
\pi_{ex} &\sim \textbf{Beta}(1,1), && (*)\\
a_n &\sim \textbf{Categorical}(1-\pi_{ex}-k_a\pi_{ex},\ \pi_{ex},\ k_a\pi_{ex}), && n\in\{1,\dots,N\},\\
\lambda_d &\sim \textbf{Gamma}(1,1), && (*)\\
\lambda_a &\sim \textbf{Gamma}(1,1), && (*)\\
\lambda_{\sigma_m}^{FRET} &\sim \textbf{Gamma}(1,1), && m\in\{1,\dots,M\}, (*)\\
w_n &\sim p(w|a_n,\lambda_d,\lambda_a,\lambda_{s_n}^{FRET}), && n\in\{1,\dots,N\},
\end{aligned}
$$

where the distribution $p(w|a_n,\lambda_d,\lambda_a,\lambda_{s_n}^{FRET})$ is the likelihood derived in Section S2 with $\boldsymbol{\rho}_{pulse}$ set by the auxiliary parameter $a_n$.

### S4 Sampling from the Posterior: Gibbs Algorithm Steps

The central object of interest in the Bayesian paradigm is the posterior

$$p(\vartheta|w_{1:N}) \propto L(w_{1:N}|\vartheta)p(\vartheta). \tag{S60}$$

where $\vartheta$ denotes the set of all unknowns including $\boldsymbol{\rho}_{start}$, rates in $\mathbf{G}_{\psi}$, and transition probabilities in $\mathbf{\Pi}_{\sigma}$. Furthermore, $p(\vartheta)$ denotes the set of priors given in Sec. S3.

In order to infer the unknown parameters, we draw numerical samples from the posterior. One way of doing this is through Markov Chain Monte Carlo (MCMC) methods, where samples from the posterior are drawn iteratively to construct a Markov chain. In this implementation, we utilize the Gibbs algorithm, where individual parameters $x$ are sampled from their conditional posterior distributions in each MCMC iteration

$$p(x|\vartheta/\{x\}, w_{1:N}), \tag{S61}$$

where $x$ is some model parameter and $\vartheta/\{x\}$ represents the set of all model parameters without $x$. In the following, we present our Gibbs sampling steps for each parameter.

#### S4.1 Photophysical rates

A photophysical rate, $\lambda$, is sampled from the conditional posterior

$$p(\lambda|\vartheta/\{\lambda\}, w_{1:N}) \propto L(w_{1:N}|\vartheta)p(\lambda), \tag{S62}$$

where prior $p(\lambda)$ is the same for all the photophysical rates

$$p(\lambda) = \mathbf{Gamma}(\lambda; 1, 1). \tag{S63}$$

This particular conditional posterior does not have a closed form, so the photophysical rates are sampled through a Metropolis-Hasting (MH) procedure. We do so by proposing new values for rates as follows

$$\lambda^* \sim \mathbf{Gamma}\left(\phi, \frac{\lambda}{\phi}\right), \tag{S64}$$

where $\phi$ is a parameter tuned to improve mixing. Subsequently, the proposal is accepted with probability given by

$$\boldsymbol{\alpha} = \min\left\{1, \frac{L(w_{1:N}|\lambda^*)\mathbf{Gamma}(\lambda^*; 1, 1)\mathbf{Gamma}(\lambda; \phi, \frac{\lambda^*}{\phi})}{L(w_{1:N}|\lambda)\mathbf{Gamma}(\lambda; 1, 1)\mathbf{Gamma}(\lambda^*; \phi, \frac{\lambda}{\phi})}\right\}, \tag{S65}$$

where $L(w_n|\lambda)$ is the likelihood for individual pulse derived in Sec. S2.

#### S4.2 Excitation Probability

To allow for direct sampling of the excitation probabilities and simplify the pulse likelihood functions derived in Sec S2, we sample the photophysical state $a_n$ immediately after the pulse. By doing so, we effectively set $\rho_{pulse}$ to be a certain photophysical state.

Since the photophysics of the individual pulses are assumed independent, we can sample each of the $a_n$ individually from their conditional posterior

$$p(a_n|\vartheta/\{a_n\}, w_{1:N}) \propto L_n(w_n|\vartheta)p(a_n) = L_n(w_n|\vartheta)\mathbf{Categorical}(a_n; \boldsymbol{\rho}_{pulse}), \tag{S66}$$

where, as derived in Sec. S2.1,

$$\boldsymbol{\rho}_{pulse} = \left(1 - \pi_{ex} - k_a\pi_{ex}, \quad \pi_{ex}, \quad k_a\pi_{ex}\right). \tag{S67}$$

Since $a_n$ represents the photophysical state immediately after the pulse, it has three photophysical states $\psi_1$, $\psi_2$, and $\psi_3$ that represent both donor and acceptor being in the ground state, the donor being excited and the acceptor being in the ground state, and the donor being in the ground state and acceptor being excited, respectively. Therefore, we sample

$$a_n \sim \mathbf{Categorical}\left(\frac{L_n(w_n|a_n = \psi_1)\xi_1}{\sum_{i=1}^{3} L_n(w_n|a_n = \psi_i)\xi_i}, \frac{L_n(w_n|a_n = \psi_2)\xi_2}{\sum_{i=1}^{3} L_n(w_n|a_n = \psi_i)\xi_i}, \frac{L_n(w_n|a_n = \psi_3)\xi_3}{\sum_{i=1}^{3} L_n(w_n|a_n = \psi_i)\xi_i}\right). \tag{S68}$$

Now, the excitation probability $\pi_{ex}$ is sampled from the conditional posterior

$$p(\pi_{ex}|\vartheta/\{\pi_{ex}\}, w_{1:N}) \propto L(w_{1:N}|\vartheta)p(\pi_{ex}) = L(w_{1:N}|\vartheta)\mathbf{Beta}(\pi_{ex}; 1, 1), \tag{S69}$$

which has likelihood-prior conjugacy because of our choice to also sample the photophysical trajectory $a_{1:N}$. Intuitively, the number of times that the photophysical trajectory records the donor being excited is the number of "successes" of a Bernoulli random variable. Therefore, we can directly sample $\pi_{ex}$ from the following probability density

$$\pi_{ex} \sim \mathbf{Beta}(1 + \sum_{i=1}^{N} \mathbb{1}\{a_i = \psi_2\}, 1 + N - \sum_{i=1}^{N} \mathbb{1}\{a_i = \psi_2\}), \tag{S70}$$

where $\sum_{i=1}^{N} \mathbb{1}\{a_i = \psi_2\}$ is the number of times that that the photophysical trajectory is in $\psi_2$.

#### S4.3 System State Trajectory

Sampling of the system state trajectory is done through a standard forward filtering backward sampling algorithm, which we briefly describe here [4]. First, we sample an initial probability vector $\boldsymbol{\pi}_0$ that is informed by the prior and the first system state of the previous trajectory.

$$\boldsymbol{\pi}_0 \sim \mathbf{Dirichlet}(\alpha((1-\kappa)\boldsymbol{\beta} + \kappa\mathbf{d}_m + \mathbf{n}_{0\cdot}), \tag{S71}$$

where $\alpha((1-\kappa)\boldsymbol{\beta} + \kappa\mathbf{d}_m$ is the prior of the transition probabilities modified with the sticky hyperparameter and $\mathbf{n}_m$ is a vector with value one at the index corresponding to $s_1$ and zero otherwise.

Next, we construct a forward filter by propagating forward using the transition matrix while taking into account observations. The first time level of the forward filter is given by,

$$\mathcal{A}_{1i} = \boldsymbol{\pi}_{0i} \times L_1(w_1|s_1 = \sigma_i), \qquad i = 1, \ldots, M_\sigma. \tag{S72}$$

This allows us to then move forward by computing for each $n$ from one to $N$,

$$\mathcal{A}_{ni} = L_n(w_n|s_1 = \sigma_i) \sum_{i=1}^{M} \boldsymbol{\pi}_{im} \mathcal{A}_{n-1,i}, \qquad i = 1, \ldots, M_\sigma,\ n = 1, \ldots, N. \tag{S73}$$

Finally, we sample the transition by recursively sampling the system state starting at the end and moving towards the first pulse in the following way

$$s_N \sim \textbf{Categorical}(\mathcal{A}_N), \tag{S74}$$

$$s_n|s_{n+1} \sim \textbf{Categorical}(\mathbf{b}_n), \tag{S75}$$

where

$$\mathbf{b}_{ni} = \frac{\boldsymbol{\pi}_{i,s_{n+1}} \mathcal{A}_{n+1,i}}{\sum_{j=1}^{M} \boldsymbol{\pi}_{j,s_{n+1}} \mathcal{A}_{n+1,j}} \qquad , i = 1, \ldots, M_\sigma,\ n = 1, \ldots, N. \tag{S76}$$

#### S4.4 Transition probabilities

The transition probabilities are sampled as vectors $\boldsymbol{\pi}_m$ that represent transition probabilities out of state $m$ from the conditional posterior

$$p(\boldsymbol{\pi}_m|\vartheta/\{\boldsymbol{\pi}_m\}, w_{1:N}) \propto L(w_{1:N}|\vartheta)p(\boldsymbol{\pi}_m) \tag{S77}$$

$$= L(w_{1:N}|\vartheta)\textbf{Dirichlet}(\boldsymbol{\pi}_m; \alpha((1-\kappa)\boldsymbol{\beta} + \kappa\mathbf{d}_m)), \tag{S78}$$

where $\alpha((1-\kappa)\boldsymbol{\beta} + \kappa\mathbf{d}_m)$ is the prior of the transition probabilities modified with the sticky hyperparameter.

These transition probability vectors are updated through likelihood-prior conjugacy with the state trajectory. Using the closed form of the conditional posterior, we sample each $\boldsymbol{\pi}_m$ through

$$\boldsymbol{\pi}_m \sim \textbf{Dirichlet}(\alpha((1-\kappa)\boldsymbol{\beta} + \kappa\mathbf{d}_m) + \mathbf{n}_{m\cdot}), \tag{S79}$$

where $\mathbf{n}_{m\cdot}$ is a vector which collects the number of each transition out of system state $\sigma_m$.

#### S4.5 Base Distribution

The sampling of the base distribution and the other hyperparameters are heavily inspired by the work of Emily Fox et. al. [5], and it is highly recommended for those interested in a more in-depth discussion of sticky iHMMs to read their work.

To sample the base distribution, we first sample auxiliary parameters $\mathbf{D}$ and $\mathbf{W}$. We start with $\mathbf{D}$. For each $i, j = 1, \ldots, M$, we set

$$M_{ij} = \sum_{k=0}^{\mathbf{n}_{ij}-1} \textbf{Bernoulli}\left(\frac{\alpha\beta_i}{j + \alpha\beta_i}\right). \tag{S80}$$

Next we sample $\mathbf{W}$, which intuitively represents the number of times a self-transition occurred because of the influence of the sticky hyperparameter.

$$W_{ii} = \textbf{Binomial}\left(M_{ii}, \frac{\kappa}{\kappa + \beta_i(1-\kappa)}\right). \tag{S81}$$

Finally, define $\overline{\mathbf{D}} = \mathbf{D} - \mathbf{W}$. We can now directly sample the base distribution through

$$\boldsymbol{\beta} \sim \textbf{Dirichlet}(\gamma\zeta + \sum_{i=1}^{M} \overline{\mathbf{D}}_{ij}), \tag{S82}$$

where $\zeta$ is an $M$ dimensional vector with all elements set to $\frac{1}{M}$.

#### S4.6 Optional Hyperparameters

Many of these sampling steps draw on auxiliary parameters $\mathbf{D}$, $\mathbf{W}$, and $\overline{\mathbf{D}}$ described in the previous section for sampling the base distribution.

##### S4.6.1 Transition probabilities concentration hyperparameter

To sample the concentration parameter $\alpha$, we sample additional auxiliary parameters, $\mathbf{r}$ and $\mathbf{s}$, which are vectors of size $M$. Define $n_{\cdot j} = \sum_{i=1}^{M} \mathbf{n}_{ij}$ as the total number of transitions into system state $\sigma_j$. We then sample

$$\mathbf{r}_i \sim \textbf{Beta}(1+\alpha, n_j), \tag{S83}$$

$$\mathbf{s}_i \sim \textbf{Bernoulli}\left(\frac{n_{\cdot j}}{n_{\cdot j} + \alpha}\right). \tag{S84}$$

We can then sample

$$\alpha \sim \textbf{Gamma}(\alpha + \sum_{i=1}^{M}\sum_{j=1}^{M} \mathbf{D}_{ij} - \sum_{i=1}^{M} \mathbf{s}, 1 - \sum_{i=1}^{M} \log(r_i)). \tag{S85}$$

##### S4.6.2 Sticky hyperparameter

Naturally the sticky hyperparameter is updated by taking the number of times self transitions occur due to the stickiness (found in $\mathbf{W}$) and using those as successes to update a Beta distribution. The sampling step is given as

$$\kappa \sim \textbf{Beta}\left(1 + \sum_{i=1}^{M} \mathbf{W}_{ii}, \phi + \sum_{i=1}^{M}\sum_{j=1}^{M} \mathbf{D}_{ij} - \sum_{i=1}^{M} \mathbf{W}_{ii}\right), \tag{S86}$$

where $\phi$ is a preset parameter that controls the "stickiness" of the HMM.

##### S4.6.3 Base distribution concentration hyperparameter

The base distribution concentrion parameter $\gamma$ requires sampling additional parameters $c$ and $p$. Additionally, define $K$ as the number of elements of $\overline{\mathbf{D}}$ that are greater than zero. Next, we sample

$$c \sim \textbf{Beta}\left(\gamma + 1, \sum_{i=1}^{M}\sum_{j=1}^{M} \overline{\mathbf{D}}_{ij}\right), \tag{S87}$$

$$p \sim \textbf{Bernoulli}\left(\frac{K}{(\sum_{i=1}^{M}\sum_{j=1}^{M} \overline{\mathbf{D}}_{ij})(1 - \log(c))}\right). \tag{S88}$$

If $p = 1$, we sample $\gamma$ as

$$\gamma \sim \textbf{Gamma}(1 + K, 1 - \log(c)), \tag{S89}$$

and otherwise we sample

$$\gamma \sim \textbf{Gamma}(K, 1 - \log(c)). \tag{S90}$$

### S5 Estimation of pre-set Parameters

Parameters associated with photon detection such as crosstalk, IRF, detection efficiency, and direct acceptor excitation are preset according to the experimental conditions. Both the label and detector quantum efficiencies are combined into $\eta_d$ and $\eta_a$. We preprocess only two parameters: IRF, by fitting IRF data to a Gaussian distribution; and background emission, which we deterimine individually.

#### S5.1 IRF

IRF data was obtained using water scattering, *i.e.*, shining the laser at a sample of water and recording the microtimes. The resulting distribution records the instrument response function, since the photons from water scattering do not experience delays due to lifetime. The expression for IRF fit using MATLAB's pre-built curve fitting tools.

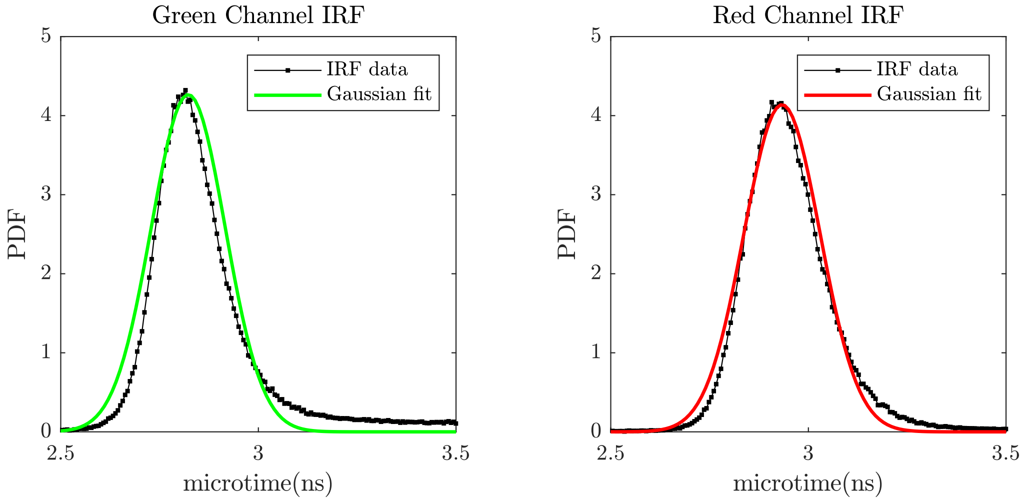


Figure S1: **IRF curve fitting.** We fit a Guassian distribution to the IRF calibration data obtained by water scattering.

#### S5.2 Background

The probability of receiving a background photon is considered constant over the course of the experiment. Let this probability be $\pi_o$. Consider a period of time with no emitting sample with pulses $1, \ldots, N$. Let $b_n$ be a label for all the pulses where $b_n = 1$ if a background photon is received in one channel and $b_n = 0$ otherwise. Given that a background photon is received, it can either come from a laser source or a uniform source. Let the probability that arrives in the laser source $\pi_b$. Let $c_k$ be a label for all background photons received with $c_k = 1$ if a photon is from the laser source, and $c_n = k$ if it is from the uniform source. In this case, we model background by

$$b_n \sim \mathbf{Bernoulli}(\pi_o), \tag{S91}$$
$$c_k | b_n = 1 \sim \mathbf{Bernoulli}(\pi_b), \tag{S92}$$
$$t_k \sim c_k \mathbf{Normal}(\mu_{IRF}, \sigma^2_{IRF}) + (1 - c_k)\mathbf{Uniform}([0, T]). \tag{S93}$$

Since the $b_n$ are known, we can directly obtain a maximum likelihood estimate for $\pi_o$. Let $K = \sum_{i=1}^{N} b_n$. Then the estimate is given by

$$\pi_o^* = \frac{K}{N}. \tag{S94}$$

We can then obtain an estimate for $\pi_b$ through

$$\pi_b^* = \arg\max_{\pi_b} \left( \prod_{i=1}^{K} \left( \pi_b \mathbf{Normal}(t_k; \mu_{IRF}, \sigma^2_{IRF}) + (1 - \pi_b)\mathbf{Uniform}([0, T]) \right) \right). \tag{S95}$$

The laser, $p_b$, and dark, $p_d$, background probabilities are then

$$p_b = \pi_0^* \pi_b^*, \tag{S96}$$
$$p_d = \pi_0^* (1 - \pi_b^*). \tag{S97}$$

An identical calculation is done for each channel.

### S6 Parameters Used for Synthetic Data Generation

Here, we detail the parameters used to produce synthetic data. Since the synthetic data algorithm incorporates crosstalk, detector efficiency, IRF, and background emissions, all of these must be set. In Table S6, the parameters not included are set according to Table S6.

| Quantity | Value Assigned | Notes |
|---|---|---|
| $\lambda_{\sigma_1 \to \sigma_2}$ | $40\,\mathrm{s}^{-1}$ | Ref. [6] |
| $\lambda_{\sigma_2 \to \sigma_1}$ | $40\,\mathrm{s}^{-1}$ | Ref. [6] |
| $\lambda_{\sigma_1}^{FRET}$ | $0.5\,\mathrm{ns}^{-1}$ | from experimental data |
| $\lambda_{\sigma_2}^{FRET}$ | $0.1\,\mathrm{ns}^{-1}$ | from experimental data |
| $\pi_{ex}$ | $5 \times 10^{-4}$ | Ref. [6] |
| $\lambda_d$ | $0.35\,\mathrm{ns}^{-1}$ | similar to ATTO 532 [7] |
| $\lambda_a$ | $0.25\,\mathrm{ns}^{-1}$ | similar to ATTO 647N [7] |
| $\mu_{IRF}$ | $2.9\,\mathrm{ns}$ | from experimental data |
| $\sigma_{IRF}^2$ | $0.001\,\mathrm{ns}^2$ | from experimental data |
| $\phi_{da}$ | 0.03 | from experimental data |
| $\phi_{ad}$ | 0.01 | from experimental data |
| $p_{bd}$ | $0.05\pi_{ex}$ | from experimental data |
| $p_{ba}$ | $0.045\pi_{ex}$ | from experimental data |
| $p_{dd}$ | $0.05\pi_{ex}$ | from experimental data |
| $p_{da}$ | $0.005\pi_{ex}$ | from experimental data |
| $\eta_d$ | 0.38 | experimental data and Ref. [7] |
| $\eta_a$ | 0.19 | experimental data and Ref. [7] |

Table S2: **Parameter values for system with two states.** Most of these values were motivated by the experimental smFRET traces gathered for this paper.

| Quantity | Value Assigned | Notes |
|---|---|---|
| $\lambda_{\sigma_1 \to \sigma_2}$ | $1200\,\mathrm{s}^{-1}$ | informed by 1 mM $MgCl_2$ HJ dynamics [8] |
| $\lambda_{\sigma_2 \to \sigma_1}$ | $1200\,\mathrm{s}^{-1}$ | informed by 1 mM $MgCl_2$ HJ dynamics [8] |
| $\lambda_{\sigma_2 \to \sigma_3}$ | $1200\,\mathrm{s}^{-1}$ | informed by 1 mM $MgCl_2$ HJ dynamics [8] |
| $\lambda_{\sigma_3 \to \sigma_2}$ | $1200\,\mathrm{s}^{-1}$ | informed by 1 mM $MgCl_2$ HJ dynamics [8] |
| $\lambda_{\sigma_1}^{FRET}$ | $0.1\,\mathrm{ns}^{-1}$ | from experimental data |
| $\lambda_{\sigma_2}^{FRET}$ | $0.4\,\mathrm{ns}^{-1}$ | from experimental data |
| $\lambda_{\sigma_3}^{FRET}$ | $0.8\,\mathrm{ns}^{-1}$ | from experimental data |
| $\pi_{ex}$ | $7.5 \times 10^{-3}$ | highest value obtained from experimental data |

Table S3: **Parameter values for system with three system states** Values that are not specified here are identical to those in Table S6 since they are set by the experimental setup and do not change from time trace to time trace.

### S7 Additional Parameter Estimates

The following figures depict the posterior distributions over all the parameters not presented in the main text.

#### S7.1 Synthetic Data with Two System States

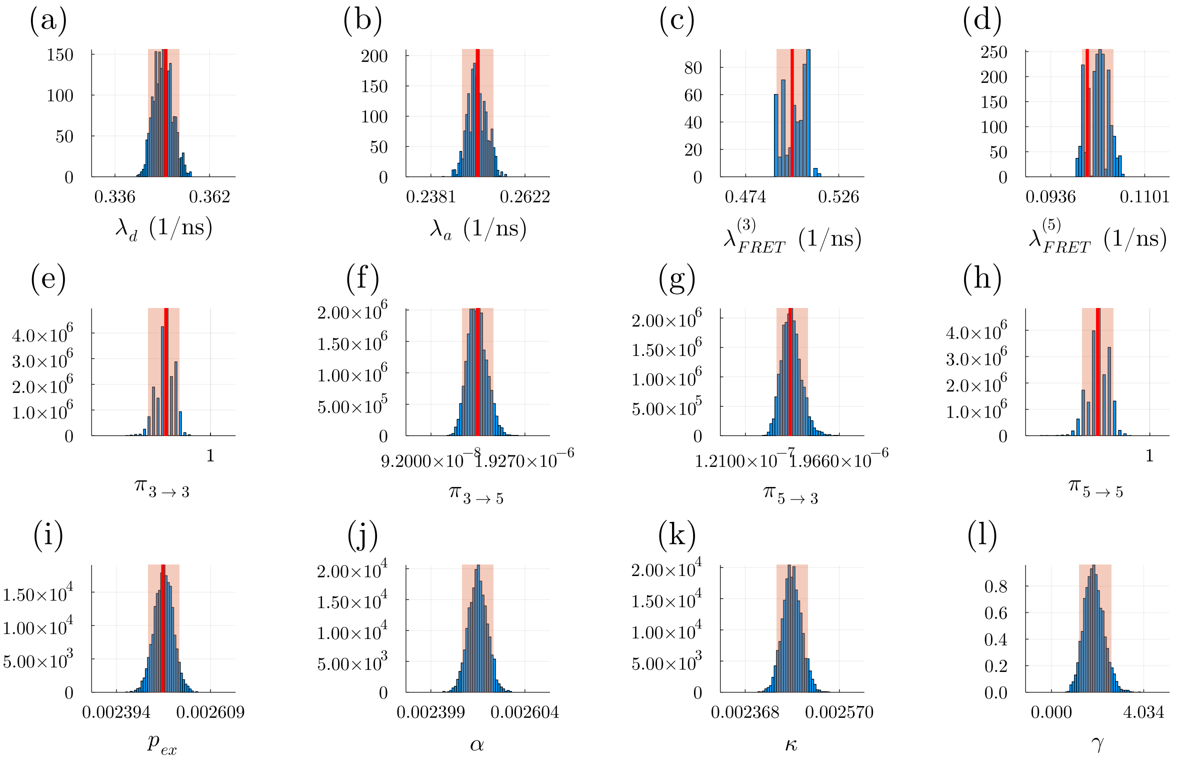


Figure S2: **Learned parameters for synthetic data with two system states.** The panels are as follows: a) donor relaxation rate; b) acceptor relaxation rate; c-d) FRET rates; e-h) system state transition probabilities for the visited states; i) excitation probability; and j-l) hyperparameters of the nonparameteric scheme, which we sample to improve mixing of the MCMC chain. The shaded regions and red lines, respectively, represent the 95% confidence interval and ground truths. The ground truth is not included for hyperparameters which are not physical quantities. The same convention is followed in the remaining figures.

#### S7.2 Synthetic Data with Three System States

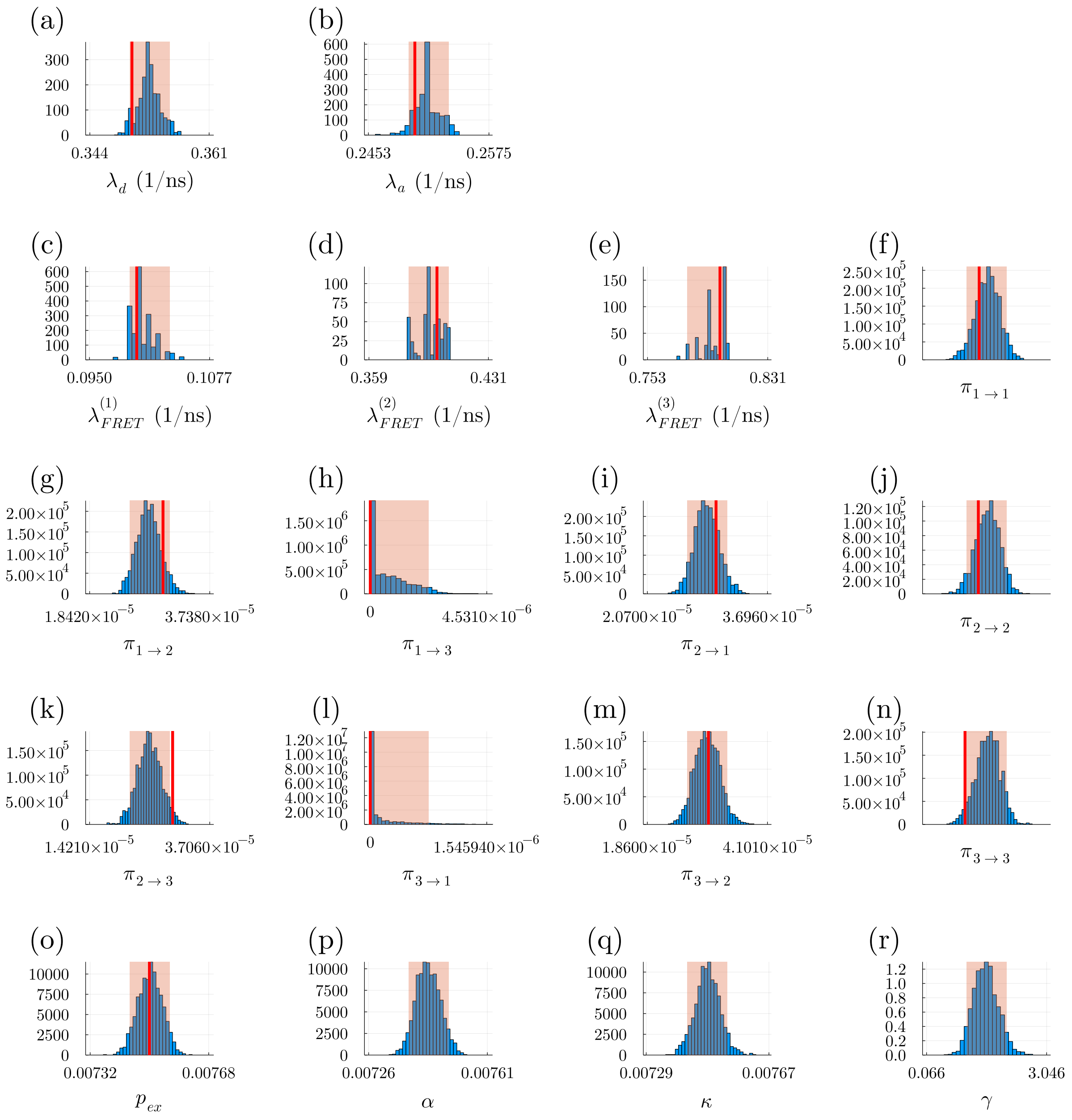


Figure S3: **Learned Parameters for Synthetic Data with three System States.** The panels are as follows: a) donor relaxation rate; b) acceptor relaxation rate; c-e) FRET rates; f-n) system state transition probabilities for the visited states; o) excitation probability; and p-r) hyperparameters of the nonparameteric scheme, which we sample for improved mixing of MCMC chain. The figure conventions are the same as those in Fig. S7.1.

#### S7.3 Experimental Data: 1 mm

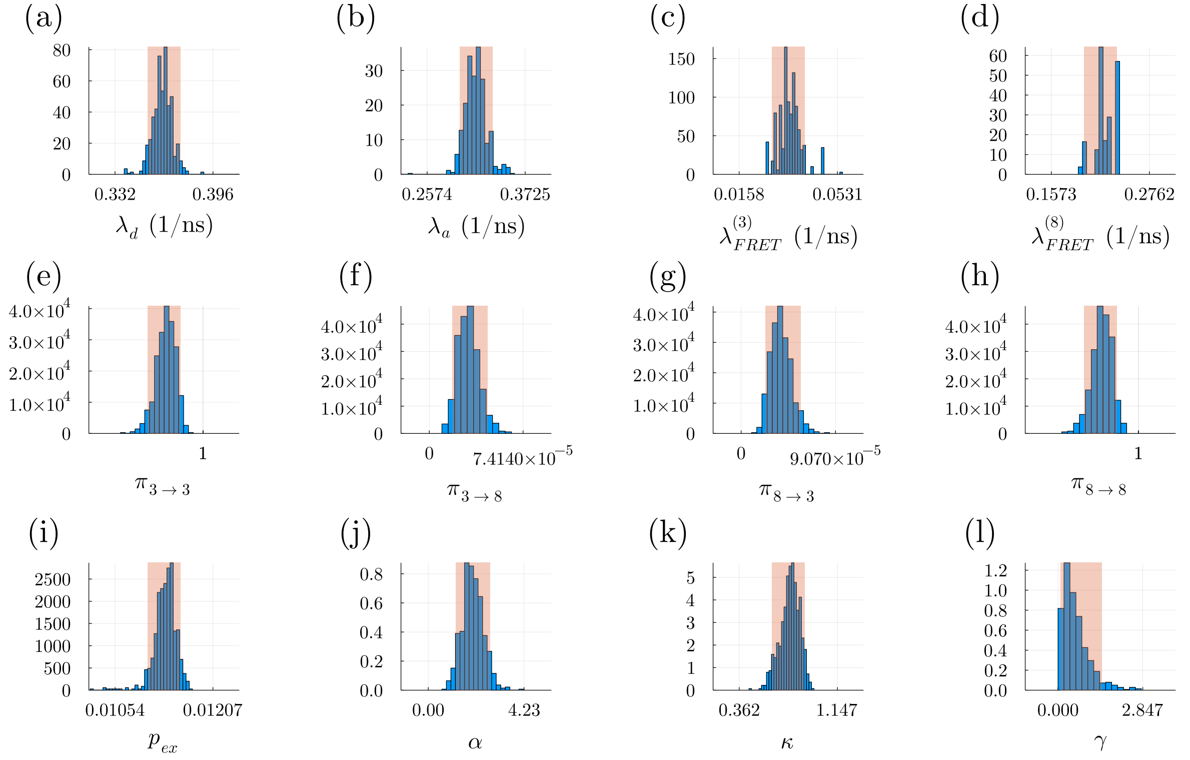


Figure S4: **Learned Parameters for 1 mm $MgCl_2$ Experimental data.** The panels are as follows: a) donor relaxation rate; b) acceptor relaxation rate; c-d) FRET rates; e-h) system state transition probabilities; i) excitation probability; and j-l) hyperparameters of the nonparametric scheme, which we sample for improved mixing of MCMC chain. The figure conventions are the same as those in Fig. S7.1.

#### S7.4 Experimental Data: 3 mm

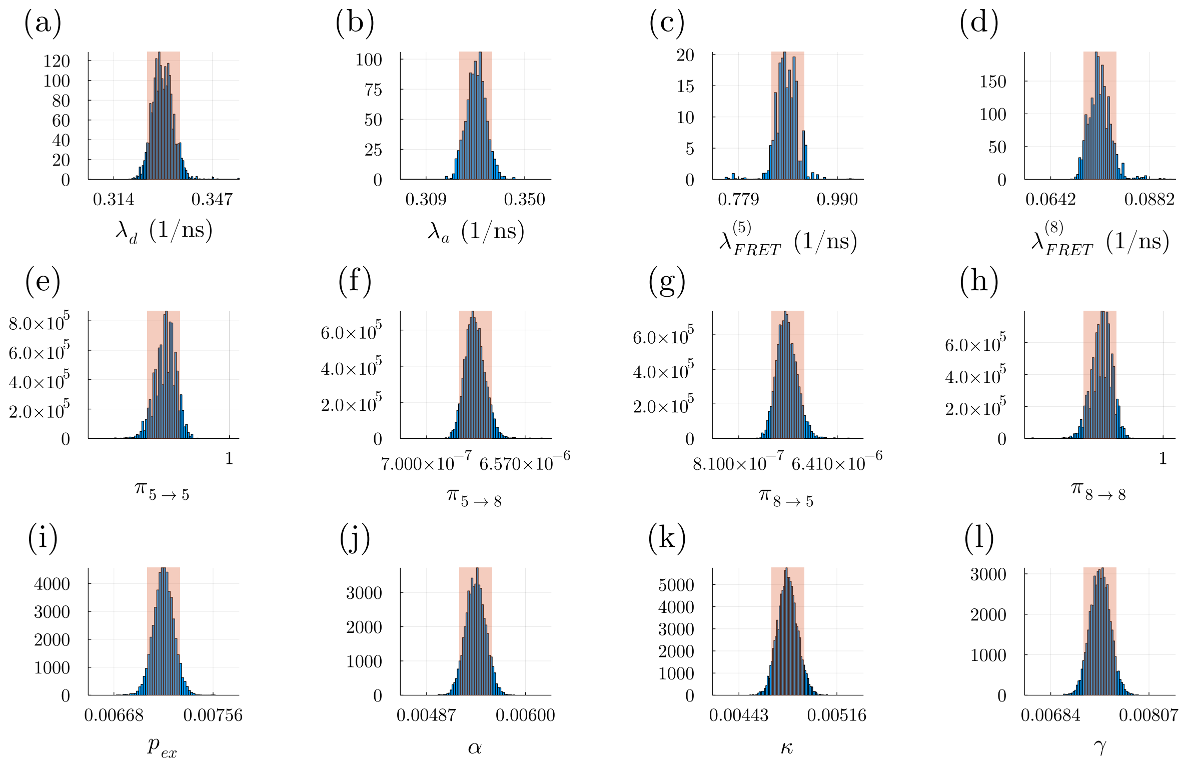


Figure S5: **Learned Parameters for 3 mm $MgCl_2$ Experimental data.** The panels are as follows: a) donor relaxation rate; b) acceptor relaxation rate; c-d) FRET rates; e)-h) system state transition probabilities for the visited states; i) excitation probability; and j-l) hyperparameters of the nonparameteric scheme, which we sample for improved mixing of MCMC chain. The figure conventions are the same as those in Fig. S7.1.

#### S7.5 Experimental Data: 5 mm

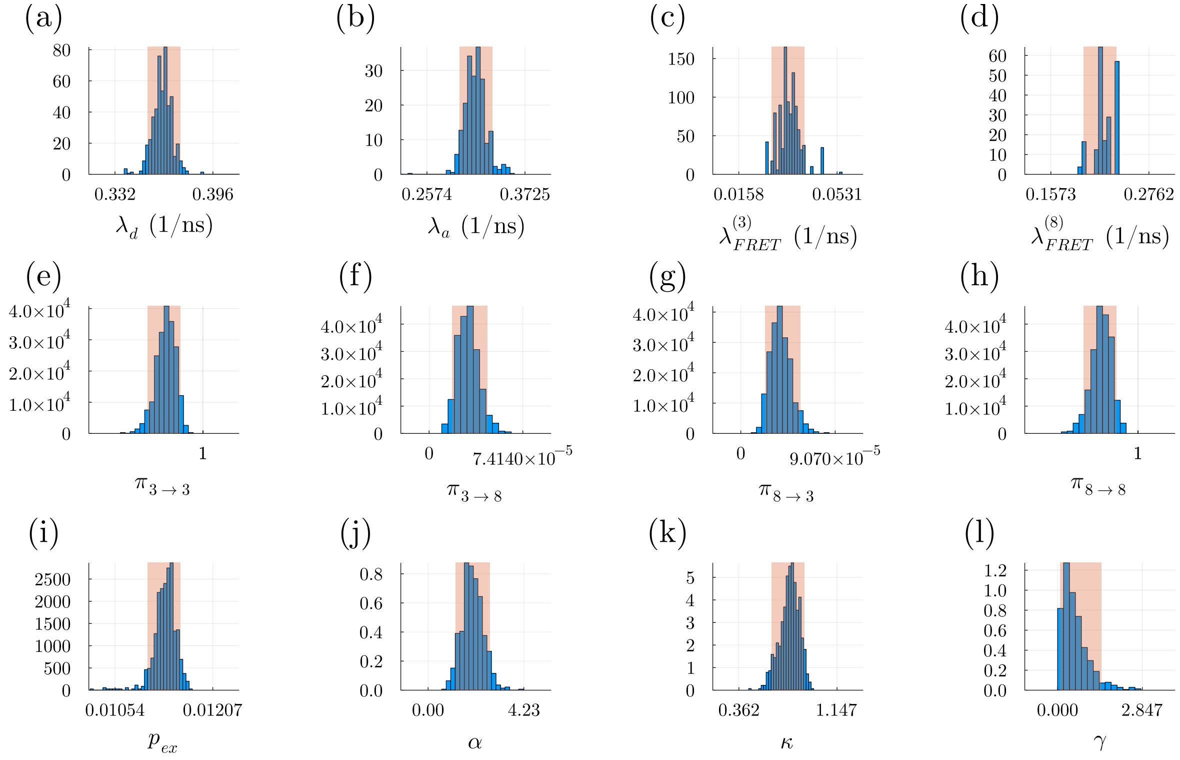


Figure S6: **Learned Parameters for 5 mm $MgCl_2$ Experimental data.** The panels are as follows: a) donor relaxation rate; b) acceptor relaxation rate; c-d) FRET rates; e-h) system state transition probabilities for the visited states; i) excitation probability; and j-l) hyperparameters of the nonparameteric scheme, which we sample for improved mixing of MCMC chain. The figure conventions are the same as those in Fig. S7.1.

#### S7.6 Experimental Data: 10 mm

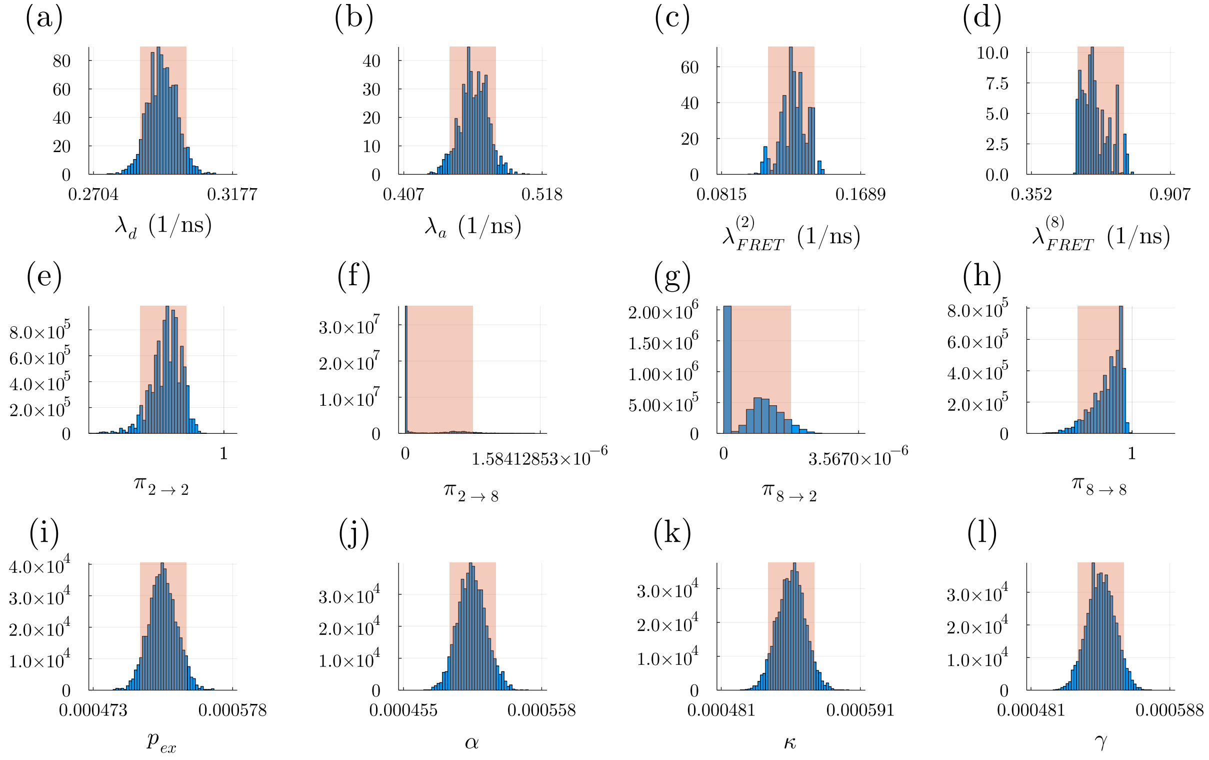


Figure S7: **Learned Parameters for 10 mm $MgCl_2$ Experimental data.** The panels are as follows: a) donor relaxation rate; b) acceptor relaxation rate; c)-d) FRET rates; e)-h) system state transition probabilities for the visited states; i) excitation probability; and j)-l) hyperparameters of the nonparameteric scheme, which we sample for improved mixing of MCMC chain. The figure conventions are the same as those in Fig. S7.1.

### Bibliography


[1] Matthew Safar, Ayush Saurabh, Bidyut Sarkar, Mohamadreza Fazel, Kunihiko Ishii, Tahei Tahara, Ioannis Sgouralis, and Steve Pressé. Single photon smFRET. III. application to pulsed illumination. *bioRxiv*, 2022.

[2] Ayush Saurabh, Matthew Safar, Ioannis Sgouralis, Mohamadreza Fazel, and Steve Pressé. Single photon smFRET. I. theory and conceptual basis. *bioRxiv*, 2022.
